# KRas, in addition to Tp53 is a driver for early carcinogenesis and a molecular target in a mouse model of invasive gastro-esophageal adenocarcinoma

**DOI:** 10.1101/2024.05.07.592904

**Authors:** Akanksha Anand, Linus Schömig, Sebastian Lange, Li Tran, Krzysztof Flisikowski, Rupert Öllinger, Roland Rad, Michael Vieth, Katja Steiger, Theresa Baumeister, Julia Strangmann, Hana Algül, Katrin Ciecielski, Katrin Böttcher, Hsin-Yu Fang, Marcos Jόse Braz Carvalho, Donja Sina Mohammad-Shahi, Sophie Gerland, Andrea Proaño-Vasco, Angelika Schnieke, Robert Thimme, Roland M. Schmid, Timothy C. Wang, Michael Quante

## Abstract

**Objective:** The incidence of gastro-esophageal adenocarcinoma (GEAC) has increased dramatically and is associated with Barrett’s Esophagus (BE). Gastric cardia progenitors are the likely origin for BE and GEAC. Here we analyze *p53, Rb1* and *Kras* alterations in Lgr5 progenitor cells during carcinogenesis.

**Design:** We introduced single and combined genetic alterations (*p53, Rb1* and *Kras*) in Lgr5-expressing progenitor cells at the inflamed gastroesophageal junction in the L2-IL1b (L2) mouse model crossed to *Lgr5-Cre^ERT^*mice. For *in-vitro* treatment we utilized mouse and human 3D organoids.

**Results:** Inactivation of *Tp53* or *Rb1* alone (L2-LP and L2-LR mice) resulted in metaplasia, and mild dysplasia, while expression of *KrasG12D* (L2-LK) accelerated dysplasia in L2-IL1b mice. Dual induction of genetic alteration in L2-LPR, L2-LKP and L2-LKR mice confirmed the accelerating role of mutant *Kras*, with the development of invasive cancer in mice with combined *Tp53* and *Kras* alteration. All three genetic events in cardia progenitor cells generated invasive cancer at 6 months of age, with chromosomal instability (CNV). The dominant role of *Kras* prompted us to treat with a SHP2 inhibitor in combination with an ERK or MEK inhibitor, leading to reduced growth in *Kras* mutant organoids. SHP2 and MEK inhibition *in-vivo* reduced *Kras* dependent tumor formation.

**Conclusion:** In the first invasive GEAC mouse model, *Kras* mutation in combination with loss of tumor suppressor genes Tp53 or Rb1 emerges as a key player in GEAC and with importance of p53 and Rb1 in promoting metaplasia. Targeting this SHP2/MEK/KRAS pathway represents a promising therapeutic option for *Kras* altered GEAC.

**What is already known on this topic:** The increased incidence of GEAC is challenging current screening and surveillance strategies. Therapeutic and preventive options are limited due to a lack of knowledge on the role of genetic alterations commonly associated with GEAC and their function during progression to dysplasia.

**What this study adds:** We generate the first invasive GEAC model and show that *KRAS* at least in combination with a second genetic alterations (*Tp53 and/or Rb1*) may be a driver of tumorigenesis, and targeting KRAS alterations could be a promising now treatment substitution.

**How this study might affect research, practice or policy:** Targeting KRAS alterations will be important for GEAC, especially as specific KRAS inhibitor are on the horizon. In addition, a concept of single genetic alteration inducing metaplasia as an adaptation to chronic inflammation might emerge as an important factor for surveillance.

## Introduction

The incidence of gastro-esophageal adenocarcinoma (GEAC), comprising both esophageal (EAC) and junctional gastric adenocarcinomas,[1] has increased in Western countries. The disparity between the large number of patients with the precursor lesions Barrett’s esophagus (BE) and the modest risk of progression to advanced GEAC poses challenges to current screening and surveillance strategies.[2] Moreover, there are limited therapeutic options for patients with GEAC. To improve both, it is important to understand the role of genetic alterations commonly associated with GEAC and their function during progression to dysplasia.

GEAC is characterized by chromosomal instability and numerous somatic alterations.[3, 4] Copy number alterations (CNA) and specific genetic mutations lead to dysregulation of key oncogenic pathways leading to clonal expansion[3, 4, 5] underlying the progression to GEAC at early BE stages.[6, 7, 8, 9, 10] Several distinct evolutionary models for genetic and phenotypic progression from BE to GEAC have been proposed; in one, an early loss of *CDKN2A* followed by loss of *TP53* function results in an increase of copy number alterations or genome doubling,[7, 8, 9] alternatively, an early loss of *TP53* followed by genome doubling and genomic instability leads to oncogene activation.[10] Loss of *CDKN2A* or *p16* is known to lead to dysregulation of *RB1*. The *TP53* and *RB1* tumor suppressor pathways both play prominent roles in blocking cancer development. Their function is inter-connected through the cyclin-dependent kinase inhibitor *p21/CDKN1A* resulting in the *TP53-p21-RB1* signaling pathway that controls transcription of a large number of genes. *KRAS* mutations are found in many human cancer types, and also reported in GEAC[11] where *KRAS* amplification is observed in 15% (Suppl. Figure 1), leading to activation of the *KRAS* oncoprotein.[12, 13]

As the vast majority of patients with gastric and BE metaplasia will never progress to GEAC,[14] identification of the mechanisms of progression could help stratify high risk patients and avoid unnecessary screening. Yet, to date, only detection of mutant *TP53* by IHC has proven a useful adjunct to confirm dysplasia. EAC may indeed derive from undifferentiated gastric progenitor cells within BE or gastric intestinal metaplasia, and it remains uncertain as to whether metaplasia itself is a necessary step in the development of GEAC, or whether there might be a shorter evolutionary trajectory from less differentiated progenitor cell types that acquire the above-mentioned genetic alterations.[15, 16] In theory, mutations that confer a survival or growth advantage within the stem cell niche will increase the succession rate and facilitate the clonal expansion and dysplastic development[1] and a dysplastic stem cell population may include a de novo gastric cancer-initiating cell responsible for neoplastic transformation[17].

In this study, we have targeted genetic alteration of *KRas, Tp53 and/or Rb1* to a non-metaplastic gastric cardia progenitor population (*Lgr5+* cells) in order accelerate carcinogenesis in the inflammatory L2-IL-1b mouse model.

## Material and methods

### Mouse strains

*L2-IL-1b (Tg(ED-L2-IL1RN/IL1B)#Tcw)*[18], *Lgr5-Cre (Lgr5-EGFP-IRES-creERT2)*[19], *Tp53 (Trp53^tm1Brn^)*[20], *Rb1 (Rb1^tm3Tyj^)*[21] and *KRasG12D (KRas^tm4Tyj^*)[22] have been described previously. *L2-IL-1b* mice were bred with *Lgr5-Cre* mice to generate L2-L (L2-IL-1b.Lgr5+/-). These mice were further crossed with *Tp53* to generate L2-LP (L2-IL-1b.Lgr5+/-.Tp53-/-), with *Rb1* to generate L2-LR (L2-IL-1b.Lgr5+/-.Rb1-/-) and with KRasG12D to generate L2-LK (L2-IL-1b.Lgr5+/-.KRas+/-). These mice were intercrossed to produce different combinations, L2-LPR (L2-IL-1b.Lgr5+/-.Tp53-/-.Rb1-/-), L2-LKP (L2-IL-1b.Lgr5+/-.KRas+/-.Tp53-/-), L2-LKR (L2-IL-1b.Lgr5+/-. KRas+/-.Rb1-/-) and L2-LKPR (L2-IL-1b.Lgr5+/-.KRas+/-Tp53-/-.Rb1-/-). Furthermore, heterozygous combinations were also generated (See Supplementary Table 1). Following weaning and genotyping, mice were fed tamoxifen food (GENOdiet CreActive T400, Radiated, Genobios Sarl, Laval, France) at the age of 4 months for 3 weeks followed by standard chow diet food (ssniff, Spezialdiäten GmbH, Soest, Germany) for another 2 months. Then the tamoxifen food was given again for 2 weeks. The mice were euthanized at the age of 7 months. All animal experiments were approved by the District Government of Upper Bavaria and performed in accordance with the German Animal Welfare and Ethical Guidelines.

## Results

### Conditional Tp53 knock-out in Lgr5+ cells accelerates proliferation and dysplasia at the Gastro-esophageal Junction in the L2-IL-1b BE mouse model

As *TP53* is altered in BE and GEAC patients,[4, 5, 23]we first characterized its role in progenitor cells at the gastro-esophageal junction (GEJ). L2-IL-1b mice[18] were crossed to *Lgr5-EGFP-IRES-creERT* (L)[19] and *Tp53* conditional knock-out mice, *Trp53tm1Brn* (P),[20] resulting in homozygous (L2-LP: *L2-IL-1b.Lgr5+/-.Tp53-/-*) or heterozygous (L2-LP+/-: *L2-IL-1b.Lgr5+/-.Tp53+/-*) mice. Compared L2-L mice (*L2-IL-1b.Lgr5+/-*) deletion of tumor suppressor *Tp53* resulted in a mildly accelerated macroscopic and histological phenotype at the age of 7 months (Figure 1A-H) with L2-LP+/- mice developing tumors of 1mm and L2-LP-/- mice with an average size of 2mm (Figure 1C). L2-LP+/- and L2-LP-/- mice showed increased metaplasia without increased infiltration of inflammatory cells (Figure 1 E, F) at the GEJ. 33% (2/6) L2-LP+/- and 20% (2/10) L2-LP mice developed low-grade dysplasia (Figure 1G, H) compared to the control L2-L group in which mice developed no dysplasia at the age of 7 months (Figure 1H). Moreover, loss of *Tp53* in progenitor cells significantly stimulated proliferation in epithelial cells at the GEJ with a 5-fold and a 6-fold increase in L2-LP+/- and L2-LP groups compared to L2-L mice (p=0.045, p=0.001; Figure I-J) along with a gradual increase in the aSMA^+^ stromal myofibroblast population surrounding the epithelial cells (Figure 1K.i). *Tp53* inactivation resulted in increased DNA damage, scored by y-H_2_AX, in L2-LP group compared to L2-L group (p=0.002; Figure 1K.ii). Knockdown of epithelial *Tp53* (Figure 1K.iii) further resulted in significantly increased p21+ epithelial cells (p=0.004; Figure 1K.iv) and stimulated epithelial apoptosis in L2-LP mice compared to controls (Figure 1K.v). These data suggest that loss of *Tp53* in epithelial progenitor cells induces hyperproliferation, p21 activation and DNA damage at the GEJ contributing to the dysplastic progression.

**Figure 1:**
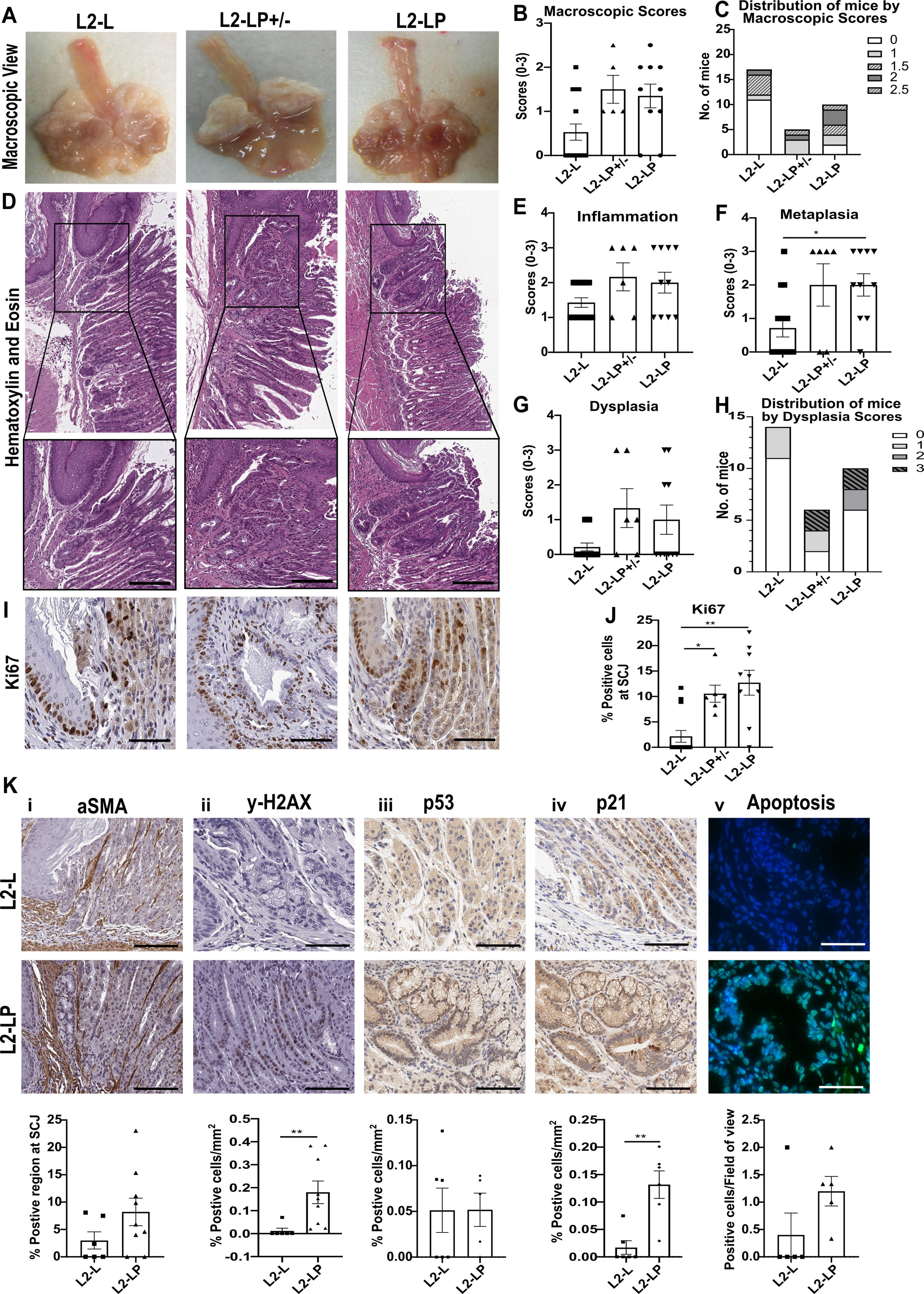
*Tp53* mutation in *Lgr5+* cells induces hyperplasia in L2-IL-1b mice. (A) Macroscopic overview of tumor formation in L2-L (*L2-IL-1b.Lgr5+/-),* L2-LP+/- (*L2-IL-1b.Lgr5+/-.Tp53+/-)*, L2-LP (*L2-IL-1b.Lgr5+/-.Tp53-/-)* mice; (B) Quantification of macroscopic scores (L2-L vs. L2-LP+/-, p=0.1216; L2-L vs. L2-LP, p=0.0608; n=5-17); (C) Distribution of mice by macroscopic scores from B; (D) Representative images of hematoxylin and eosin staining of L2-L, L2-LP+/- and L2-LP mice to assess the histopathology, scale bars 100μm and 200μm; (E) Inflammation (L2-L vs. L2-LP+/-, p=0.2720; L2-L vs. L2-LP, p=0.3779; n=6-14); (F) Metaplasia (L2-L vs. L2-LP+/-, p=0.1278; L2-L vs. L2-LP, *p=0.0444; n=6-14); (G) Dysplasia (L2-L vs. L2-LP+/-, p=0.1193; L2-L vs. L2-LP, p=0.5319; n=6-14); (H) Distribution of mice by dysplasia scores from G; (I) Representative images of immunohistochemistry for Ki67 for L2-L, L2-LP+/- and L2-LP mice, scale bar 50μm; (J) Immunohistochemistry for Ki67 (L2-L vs. L2-LP+/-, *p=0.0455; L2-L vs. L2-LP, **p=0.0011; n=6-14); (K) Panel of tumor microenvironment and DNA damage comparing L2-L and L2-LP cohorts: i. Immunohistochemistry for a-SMA (p=0.1728; n=6-8), ii. Immunohistochemistry for y-H_2_AX (**p=0.0022; n=6-8), iii. Immunohistochemistry for Tp53 (p=0.9177; n=5-6), iv. Immunohistochemistry for p21 (**p=0.0043; n=6), v. TUNEL assay for apoptosis (p=0.0794; n=6). Scale bar for K.i-v 50μm. For statistical analysis, B,E,F,G,J: Kruskal-Wallis test was performed in addition to adjusted p-value with Dunn’s multiple comparison test; K.i-v: Mann-Whitney test was performed.

### Conditional inactivation of Rb1 in Lgr5+ cells induces low-grade dysplasia in L2-IL-1b mice

As *Tp53* inactivation was not found to be sufficient for the progression to cancer in L2-IL-1b mice, we next examined the effect of another cell cycle regulator, *RB1*, which is located downstream of its *CDKN2A* effectors *p14^ARF^* and *p16^INK4A^.* We utilized *Rb1tm3Tyj* (R) mice)[21] in combination with L2-L mice to produce conditional homozygous (L2-LR*: L2-IL-1b.Lgr5+/-.Rb1-/-*) and heterozygous (L2-LR+/-: *L2-IL-1b.Lgr5+/-.Rb1+/-*) mice. *Rb1* inactivation in *Lgr5+* cells in 7-month-old mice led to formation of scattered lesions in the GEJ, reflecting an increase in tumor burden in L2-LR+/- and L2-LR compared to L2-L mice (Figure 2A-B). L2-LR+/- mostly developed lesions of <1mm compared to L2-LR which developed lesions of 1-2mm (Figure 2C). Rb1 inactivation resulted in significantly increased inflammation and extended metaplasia at the GEJ in L2-LR+/- and L2-LR compared to L2-L mice (Figure 2D-F), which progressed to high-grade dysplasia in 25% (2/8) L2-LR+/- mice and low-grade dysplasia in 27% (3/11) L2-LR mice (Figure 2G-H). Rb1 inactivation led to higher proliferation of epithelial cells at GEJ (p=0.015, p=0.004; Figure 2I-J) along with an increase in myofibroblasts (Figure 2K.i), and DNA damage (p=0.015; Figure 2K.ii). Of note, Rb1 inactivation resulted in higher epithelial Tp53 (Figure 2K.iii) and p21 expressions compared to L2-L mice (p=0.004; Figure 2K.iv) correlating with increased cell death (Figure 2K.v). In summary, *Rb1* elimination only mildly accelerated dysplasia in the epithelial cells but in distinctively increase metaplasia.

**Figure 2:**
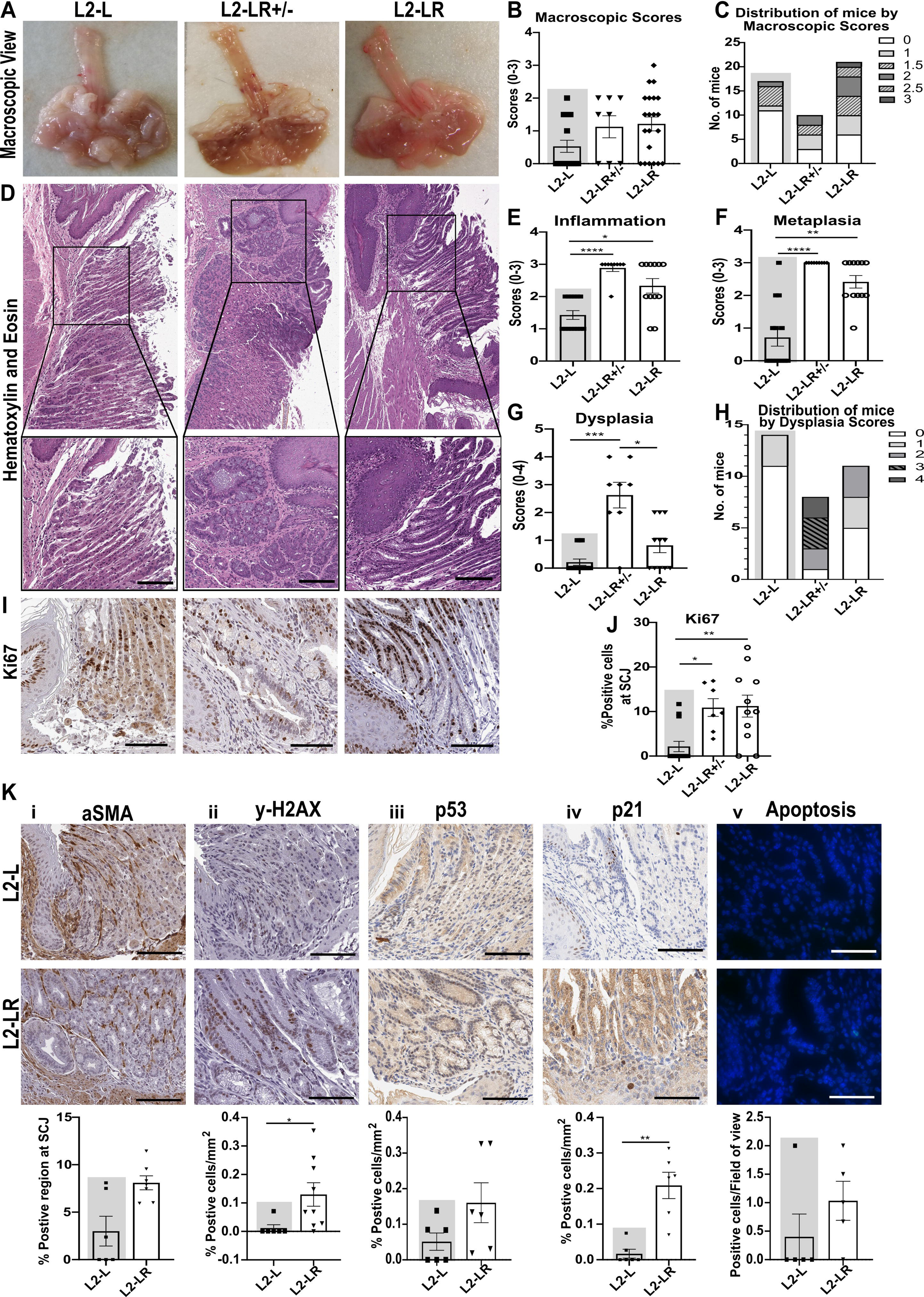
*Rb1* inactivation in *Lgr5+* mice induces low-grade dysplasia in L2-IL-1b mice. (A) Macroscopic overview of tumor formation in L2-L (*L2-IL-1b.Lgr5+/-),* L2-LR+/- (*L2-IL-1b.Lgr5+/-.Rb1+/-)*, L2-LR (*L2-IL-1b.Lgr5+/-.Rb1-/-)* mice; (B) Quantification of macroscopic scores (L2-L vs. L2-LR+/-, p=0.3345; L2-L vs. L2-LR, p=0.0695; n=8-21); (C) Distribution of mice by macroscopic scores from B; (D) Representative images of hematoxylin and eosin staining of L2-L, L2-LR+/- and L2-LR mice to assess the histopathology, scale bars 100μm and 200μm; (E) Inflammation (L2-L vs. L2-LR+/-, ****p<0.0001; L2-L vs. L2-LR, *p=0.0167; n=9-14); (F) Metaplasia (L2-L vs. L2-LR+/-, ****p<0.0001; L2-L vs. L2-LR, **p=0.0038; n=9-14); (G) Dysplasia (L2-L vs. L2-LR+/-, ***p=0.0003; L2-L vs. L2-LR, p=0.0413; n=9-14); (H) Distribution of mice by dysplasia scores from G; (I) Representative images of immunohistochemistry for Ki67 for L2-L, L2-LR+/- and L2-LR mice, scale bar 50μm; (J) Immunohistochemistry for Ki67 (L2-L vs. L2-LR+/-, *p=0.0149; L2-L vs. L2-LR, **p=0.0044; n=7-14); (K) Panel of tumor microenvironment and DNA damage comparing L2-L and L2-LR cohorts: i. Immunohistochemistry for a-SMA (p=0.0501; n=6-7), ii. Immunohistochemistry for y-H_2_AX (*p=0.0146; n=6-9), iii. Immunohistochemistry for Tp53 (p=0.1277; n=6), iv. Immunohistochemistry for p21 (**p=0.0043; n=6), v. TUNEL assay for apoptosis (p=0.2063; n=5). Scale bar for K.i-v 50μm. For statistical analysis, B,E,F,G,J: Kruskal-Wallis test was performed in addition to adjusted p-value with Dunn’s multiple comparison test; K.i-v: Mann-Whitney test was performed. Groups with grey backgrounds are the same mice cohorts as their respective groups in Figure 1.

### Combined elimination of *Tp53* and *Rb1* in *Lgr5+* cells cause BE to progress to high-grade dysplasia

Next, we tested a combined inactivation of *Tp53* and *Rb1* in *Lgr5+* cells. Thus, we crossed L2-LP and L2-LR to generate the following combinations: L2-LP+/-R+/- *(L2-IL-1b.Lgr5+/-.Tp53+/-.Rb1+/-*); L2-LP+/-R-/- *(L2- IL-1b.Lgr5+/-.Tp53+/-.Rb1-/-*); L2-LP-/-R+/- *(L2-IL-1b.Lgr5+/-.Tp53-/-.Rb1+/-*) and L2-LPR *(L2-IL-1b.Lgr5+/-.Tp53-/-.Rb1-/-*) mice. Co-inactivation of *Tp53* and *Rb1* (L2-LPR mice) accelerated disease progression but did not yet induce invasive cancer. Assessment of macroscopic tumor burden demonstrated that only 7% (1/14) of L2-LPR mice developed lesions >2mm (Figure 3A-C). L2-LP+/-R+/-, L2-LP+/-R-/- and L2-LP-/-R+/- mice revealed increased inflammation and metaplasia at the GEJ compared to L2-L mice, and in L2-LPR mice this increase was significant (p=0.04, 0.002; Figure 3D-F), again suggesting that the inactivation of *Rb1* even with loss of *Tp53* induced a metaplastic phenotype rather than a dysplastic progression (Suppl. Figure 2A-B). Of note, 20% (2/10) L2-LPR mice developed high-grade dysplasia compared to none in the L2-L group. Moreover, we found a significantly higher proliferation (p=0.012; Figure 3I-J) and increased stromal myofibroblast expression in L2-LPR compared to L2-L mice (Figure 3K.i). DNA damage in epithelial cells from L2-LPR mice was also significantly increased compared to L2-L mice (p=0.002; Figure 3K.ii), correlating with a change in *Tp53* in the tissue (Figure 3K.iii) and an increase in p21 expression (p=0.013; Figure 3K.iv) resulting in increased cell death (p=0.024; Figure 3K.v).

**Figure 3:**
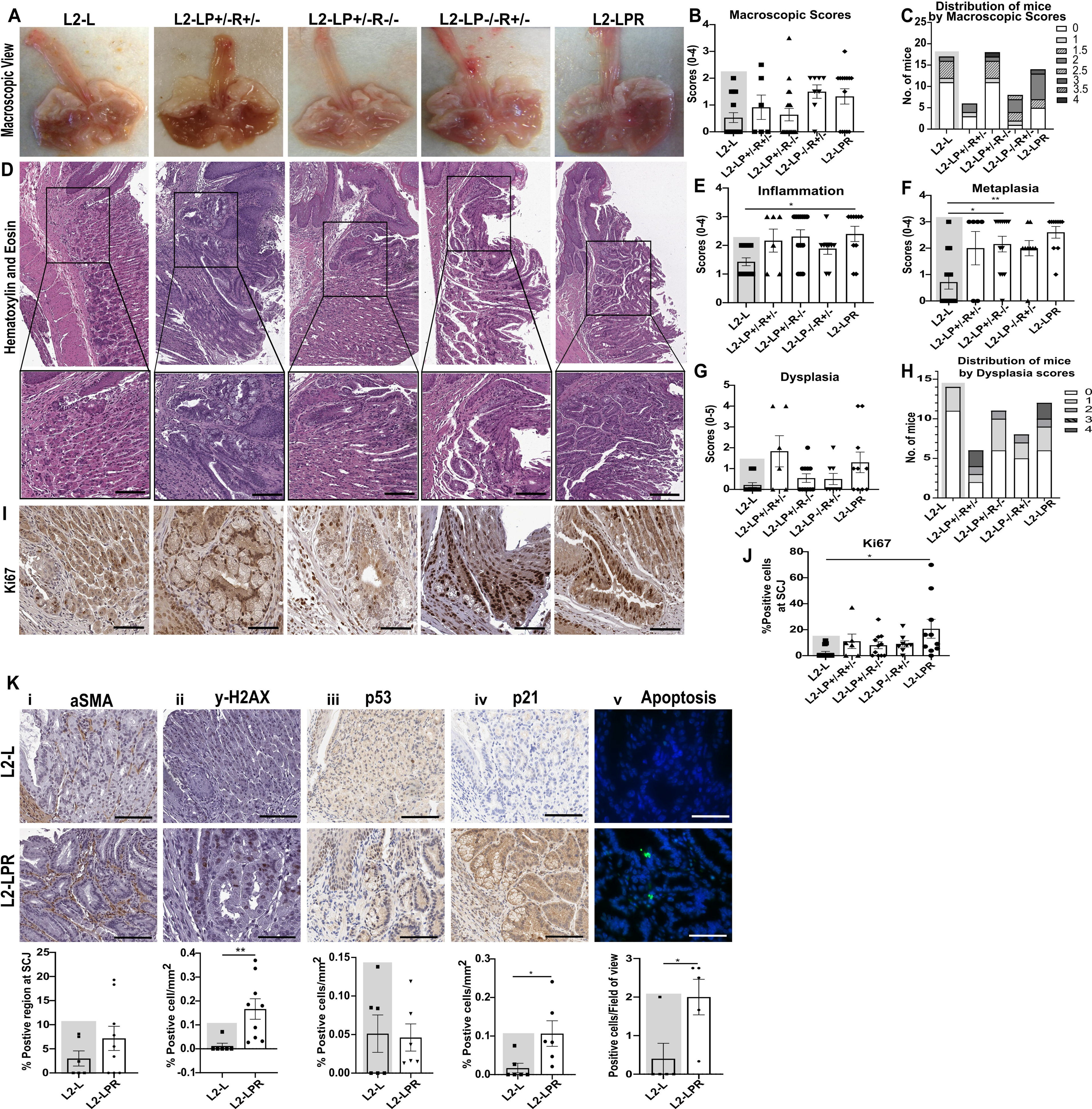
Co-inactivation of *Tp53* and *Rb1* induces high-grade dysplasia in L2-IL-1b mice. (A) Macroscopic overview of tumor formation in L2-L (*L2-IL-1b.Lgr5+/-),* L2-LP+/-R+/- (*L2-IL-1b.Lgr5+/-.Tp53+/-.Rb1+/-)*, L2-LP+/-R-/- (*L2-IL-1b.Lgr5+/-.p5+/-.Rb1-/-)*, L2-LP-/-R+/- (*L2-IL-1b.Lgr5+/-.p5-/-.Rb1+/-)*, L2-LPR (*L2-IL-1b.Lgr5+/-.Tp53-/-.Rb1-/-)* mice; (B) Quantification of macroscopic scores (L2-L vs. L2-LPR, p=0.1868; n=6-18); (C) Distribution of mice by macroscopic scores from B; (D) Representative images of hematoxylin and eosin staining of L2-L, L2-LP+/-R+/-, L2-LP+/-R-/-, L2-LP-/-R+/-, L2-LPR mice to assess the histopathology, scale bars 100μm and 200μm; (E) Inflammation (L2-L vs. L2-LPR, *p=0.0406; n=6-14); (F) Metaplasia (L2-L vs. L2-LP+/-R-/-, *p=0.0231; L2-L vs. L2-LPR, **p=0.0019; n=6-14); (G) Dysplasia (L2-L vs. L2-LPR, p=0.3272; n=6-14); (H) Distribution of mice by dysplasia scores from G; (I) Representative images of immunohistochemistry for Ki67 for L2-L, L2-LP+/-R+/-, L2-LP+/-R-/-, L2-LP-/-R+/-, L2-LPR mice, scale bar 50μm; (J) Immunohistochemistry for Ki67 (L2-L vs. L2-LPR, *p=0.0123; n=6-14); (K) Panel of tumor microenvironment and DNA damage comparing L2-L and L2-LPR cohorts: i. Immunohistochemistry for a-SMA (p=0.3267; n=6-9), ii. Immunohistochemistry for y-H_2_AX (**p=0.0022; n=6-9), iii. Immunohistochemistry for Tp53 (p=0.8139; n=6), iv. Immunohistochemistry for p21 (*p=0.0130; n=6), v. TUNEL assay for apoptosis (*p=0.0238; n=5). Scale bar for K.i-v 50μm. For statistical analysis, B,E,F,G,J: Kruskal-Wallis test was performed in addition to adjusted p-value with Dunn’s multiple comparison test; K.i-v: Mann-Whitney test was performed. Groups with grey backgrounds are the same mice cohorts as their respective groups in Figure 1.

### *KrasG12D* mutation emerged as a driver for progression from BE to EAC in the L2-IL-1b mouse model

Due to long latency and low penetrance of cancer development by co-deletion of *Tp53* and *Rb1* in *Lgr5+* cells at the age of 7 months, we introduced a third genetic alteration, *KrasG12D*, utilizing *Krastm4Tyj* (K) mice,[22] generating a *Kras*-specific mouse model L2-LK (*L2-IL-1b.Lgr5+/-.Kras+/-*). We further generated double transgenic mouse models with *Tp53* and *Rb1* inactivation, which led to homozygous combinations L2-LKP (*L2-IL-1b.Lgr5+/-.Kras+/-.Tp53-/-*) and L2-LKR (*L2-IL-1b.Lgr5+/-. Kras+/-.Rb1-/-*) and their heterozygous counterparts L2-LKP+/- (*L2-IL-1b.Lgr5+/-. Kras+/-.Tp53+/-*) and L2-KR+/- (*L2-IL-1b.Lgr5+/-. Kras+/-.Rb1+/-*). Finally, we combined all the genetic alterations, generating a triple transgenic epithelial progenitor specific model resulting in L2-LKPR mice (*L2-IL-1b.Lgr5+/-.Kras+/-Tp53-/-.Rb1-/-*) including its different heterozygous combinations: L2-LKP+/-R+/- (*L2-IL-1b.Lgr5+/-.Kras+/-Tp53+/-.Rb1+/-*); L2-LKP+/- R-/- (*L2-IL-1b.Lgr5+/-.Kras+/-Tp53+/-.Rb1-/-*); L2-LKP-/-R+/- (*L2-IL-1b.Lgr5+/-.Kras+/-Tp53-/-.Rb1+/-*) mice.

The conditional *KrasG12D* mutation alone in *Lgr5+* GEJ cells significantly accelerated dysplasia (Figure 4A, D) by the age of 7 months. Combinations of *KrasG12D* with *Rb1* and *Tp53* inactivation (L2-LKR and L2-LKP mice) resulted in the development of invasive cancer at the GEJ (Figure 4A, D). Cancer development was accelerated even further in L2-LKPR mice (Figure 4A, D), wherein invasive cancer developed at 6 months of age. We observed a significant increase in macroscopic tumor burden in L2-LKR and L2-LKP mice, forming tumors >5mm in 19% (3/16) L2-LKR mice, in 33% (2/6) L2-LKP mice, and 41% (7/17) in L2-LKPR compared to L2-L mice (p<0.0001; Figure 4A-C). We also observed increased tumor formation in L2-LKP+/-R-/- and L2-LKP-/-R+/- mice, suggesting that in addition to the *Kras* mutation the inactivation of at least one tumor suppressor gene is required, to induce tumor growth (Suppl. Figure 3A-C). While 30% (3/10) of L2-LK mice developed high-grade dysplasia, 64% (7/11) of L2-LKR, 50% (6/12) of L2-LKP and 81% (13/16) of L2-LKPR mice developed invasive cancer (Figure 4D-H). We observed cancer formation in L2-LKP+/-R+/-, L2-LKP+/-R-/- and L2-LKP-/-R+/- mice (Suppl. Figure 3D-H) supported by increased proliferation (Suppl. Figure 3I-J). To score cancer malignancy tumor grading equivalent to the human setting was performed. G1-G2 tumors were present in L2-LKR mice, whereas L2-LKP and L2-LKPR mice developed up to G3 grade tumors (Figure 4I). Tumors showed excessive hyperproliferation in double and triple mutant mice, with significantly higher proliferative epithelial cell numbers in L2-LKR (p=0.0005), L2-LKP (p=0.0008) and L2-LKPR (p<0.0001) mice compared to L2-L mice (Figure 4J-K).

**Figure 4:**
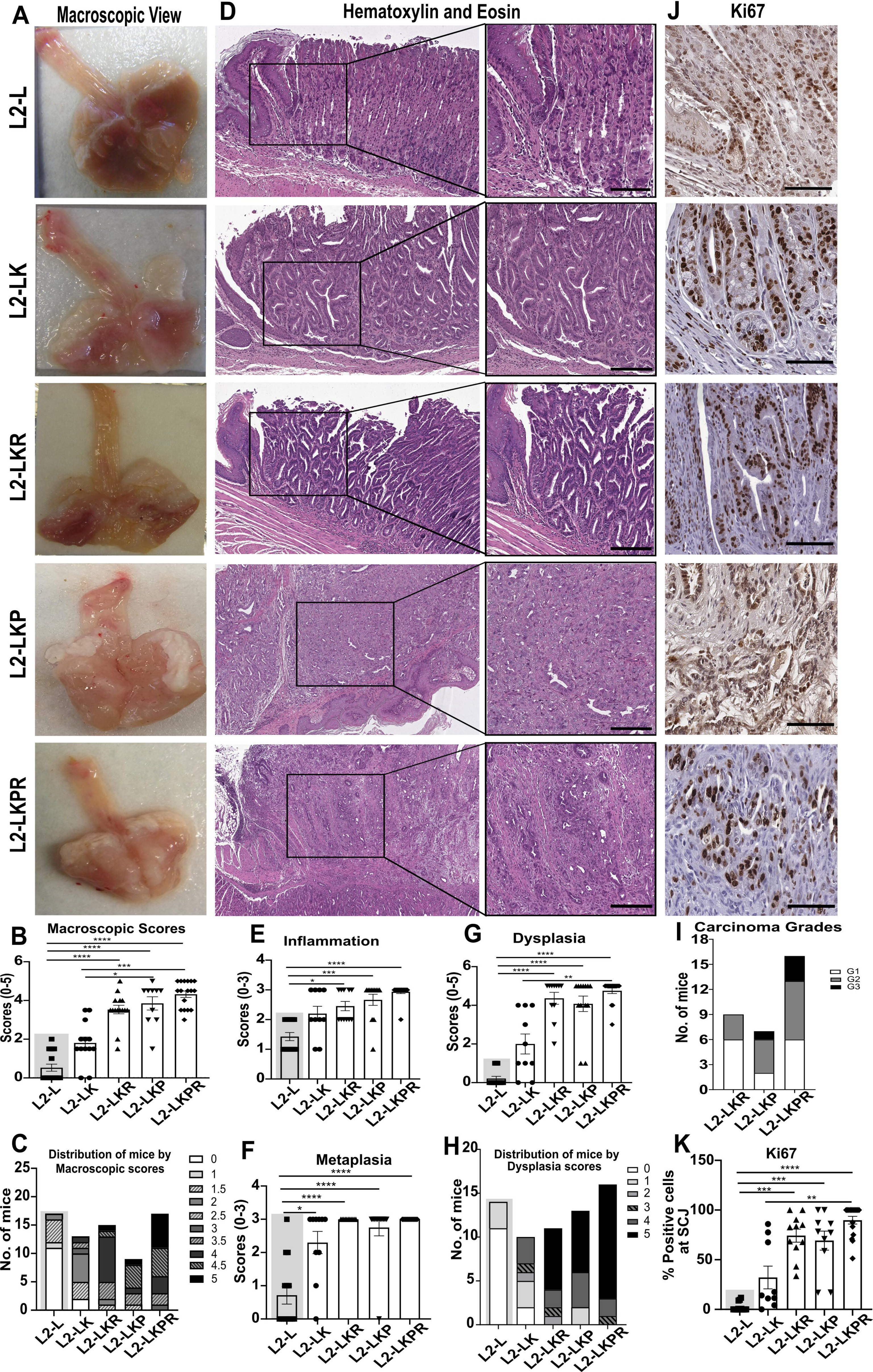
*KrasG12D* alteration in combination with *Tp53* and *Rb1* induces invasive cancer in L2-IL-1b mice. (A) Macroscopic overview of tumor formation in L2-L (*L2-IL-1b.Lgr5+/-),* L2-LK (*L2-IL-1b.Lgr5+/-.Kras+/-)*, L2-LKR (*L2-IL-1b.Lgr5+/-.Kras+/-.Rb1-/-)*, L2-LKP (*L2-IL-1b.Lgr5+/-.Kras+/-Tp53-/-)*, L2-LKPR (*L2-IL-1b.Lgr5+/-.Kras+/-.Tp53-/-.Rb1-/-)* mice; (B) Quantification of macroscopic scores (L2-L vs. L2-LK, p>0.9999; L2-L vs. L2-LKR, ****p<0.0001; L2-L vs. L2-LKP, ****p<0.0001; L2-L vs. L2-LKPR, ****p<0.0001,L2-LK vs. L2-LKP, *p=0.0426; L2-LK vs. L2-LKPR, ***p=0.0002; n=10-17); (C) Distribution of mice by macroscopic scores from B; (D) Representative images of hematoxylin and eosin staining of L2-L, L2-LK, L2-LKR, L2-LKP, L2-LKPR mice to assess the histopathology, scale bars 100μm and 200μm; (E) Inflammation (L2-L vs. L2-LK, p=0.2041; L2-L vs. L2-LKR, *p=0.0235; L2-L vs. L2-LKP, ***p=0.0004; L2-L vs. L2-LKPR, ****p<0.0001; n=10-16); (F) Metaplasia (L2-L vs. L2-LK, *p=0.0205; L2-L vs. L2-LKR, ****p<0.0001; L2-L vs. L2-LKP, ****p<0.0001; L2-L vs. L2-LKPR, ****p<0.0001; n=10-16); (G) Dysplasia (L2-L vs. L2-LK, p>0.9999; L2-L vs. L2-LKR, ****p<0.0001; L2-L vs. L2-LKP, ****p<0.0001; L2-L vs. L2-LKPR, ****p<0.0001; n=10-16); (H) Distribution of mice by dysplasia scores from G; (I) Representative images of immunohistochemistry for Ki67 for L2-L, L2-LK, L2-LKR, L2-LKP, L2-LKPR mice, scale bar 50μm; (J) Immunohistochemistry for Ki67 (L2-L vs. L2-LK, p=0.6351; L2-L vs. L2-LKR, ***p=0.0005; L2-L vs. L2-LKP, ****p=0.0008; L2-L vs. L2-LKPR, ****p<0.0001; L2-LK vs. L2-LKPR, **p=0.0080, n=9-16); For statistical analysis, Kruskal-Wallis test was performed in addition to adjusted p-value with Dunn’s multiple comparison test. Groups with grey backgrounds are the same mice cohorts as their respective groups in Figure 1.

Tumor formation was accompanied by an increase in the stromal microenvironment, with a significant increase in stromal myofibroblasts in L2-LKPR compared to L2-L mice (Figure 5A-B). *Kras* mutated mice also showed increased DNA damage in L2-LK, L2-LKR (p=0.027), L2-LKP (p=0.004) and L2-LKPR (p=0.0155) compared to L2-L mice (Figure 5C-D). This correlated with higher expression of Tp53 and p21 in L2-LK mice. L2-LKR, L2-LKP and L2-LKPR mice showed a lower Tp53 and also p21 expression in the tissue. Increased epithelial apoptosis was observed in in L2-LK, L2-LKP and L2-LKPR mice (Figure 5I-J). These data suggest a pivotal function of altered KRas during disease progression to EAC. However, the additional inactivation of at least one tumor suppressor genes was needed to induce invasive cancer.

**Figure 5:**
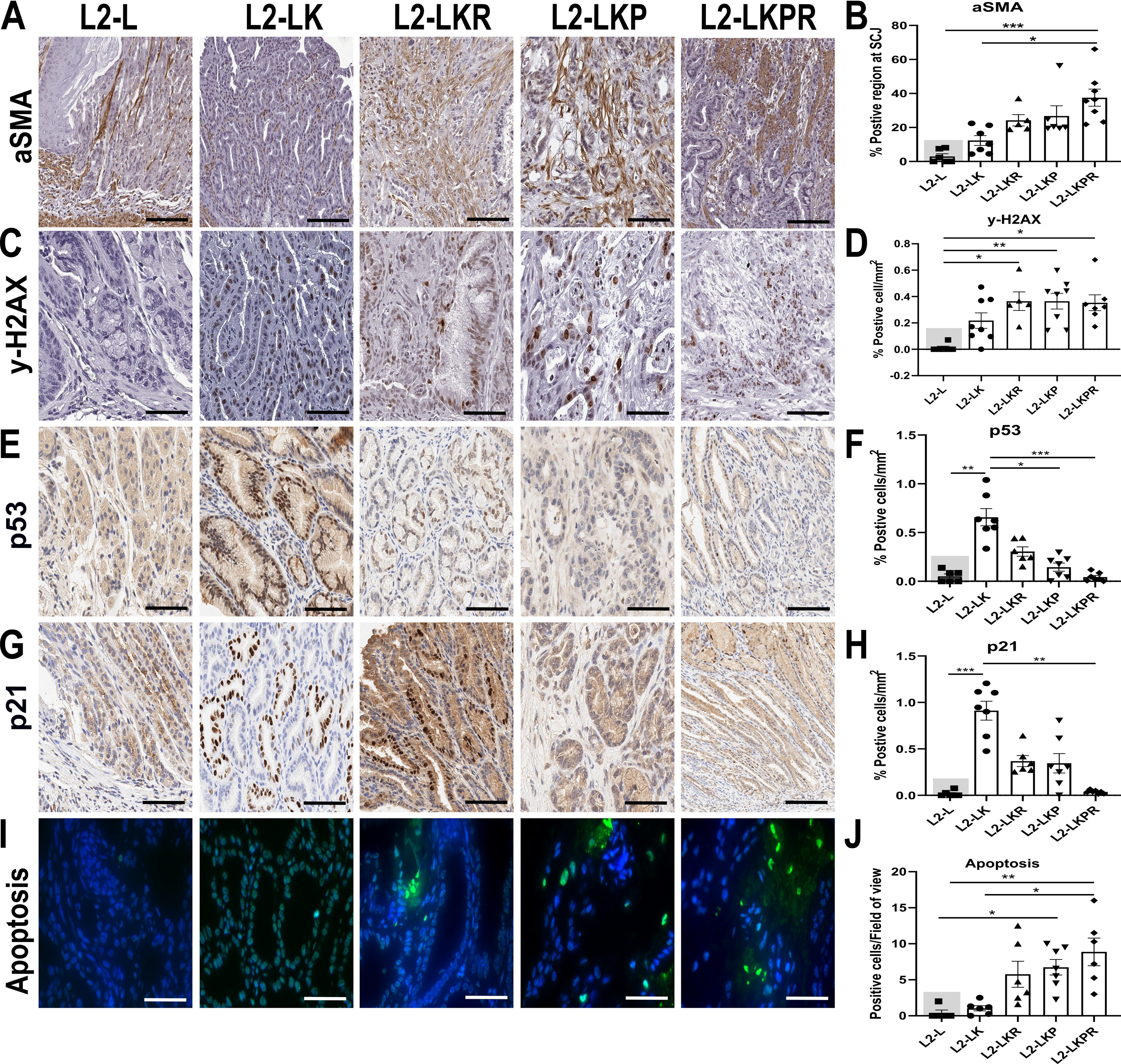
*KrasG12D* mutation induces changes in the epithelial and the stromal cells of the tumor microenvironment. Panel of tumor microenvironment and DNA damage comparing L2-L mice with different *Kras*-mutated cohorts: (A) Representative images and (B) Immunohistochemistry for a-SMA (L2-L vs. L2-LKPR, ***p=0.0002; L2-LK vs. L2-LKPR, *p=0.0170; n=5-8), (C) Representative images and (D) Immunohistochemistry for y-H_2_AX (L2-L vs. L2-LKR, *p=0.0273; L2-L vs. L2-LKP, **p=0.0042; L2-L vs. L2-LKPR, *p=0.0155; n=5-8), (E) Representative images and (F) Immunohistochemistry for Tp53 (L2-L vs. L2-LK, ***p=0.0007; L2-L vs. L2-LKP, **p=0.0042; L2-LK vs. L2-LKP, *p=0.0365; L2-LK vs. L2-LKPR, *p=0.0006; n=6-7), (G) Representative images and (H) Immunohistochemistry for p21 (L2-L vs. L2-LK, ***p=0.0001; L2-LK vs. L2-LKPR, **p=0.0015; n=6-7), (I) Representative images and (J) TUNEL assay for apoptosis (L2-L vs. L2-LKR, p=0.0816; L2-L vs. L2-LKP, *p=0.0218; L2-L vs. L2-LKPR, **p=0.0038; L2-LK vs. L2-LKPR, *p=0.0189; n=5-7). Scale bar 50μm for all the images. For statistical analysis, B,D,F,H,J: Kruskal-Wallis test was performed in addition to adjusted p-value with Dunn’s multiple comparison test; Groups with grey backgrounds are the same mice cohorts as their respective groups in Figure 1.

### *KrasG12D* altered epithelial cells exhibit distinct features in in-vitro organoid cultures

To understand the specificity of these genetic alterations in epithelial cells, we utilized the 3D organoid model system. Organoids isolated from *KrasG12D* mice, i.e., L2-LK, L2-LKR, L2-LKP and L2-LKPR, developed a distinct budding formation on the boundary of organoids (Figure 6A, indicated in red arrows), which was not observed in control, L2-LR, L2-LP and L2-LPR organoids. Budding can be suggestive of increased proliferation as previously observed in intestinal epithelial organoids[24]. Histopathological analysis showed increased apoptotic and atypical cellular formation in L2-LK, L2-LKR, L2-LKP and L2-LKPR organoids compared to more columnar cells states in L2-L, L2-LR, L2-LP and L2-LPR groups (Figure 6B). Organoids with mutant *Kras* showed higher proliferation compared to control (Figure 6C-D). Furthermore, mutant *Kras* affected the survival of the organoids. While L2-L, L2-LR, L2-LP and L2-LPR organoids could be maintained for maximum five passages, L2-LK organoids survived more than 5 passages, L2-LKR and L2-LKP organoids up to 10, and L2-LKPR for more than 15 passages (Figure 6E). These findings were in concordance with the histopathological analysis, where we graded the organoids based on the cellular architecture. Compared to L2-L, organoids from L2-LKR (p=0.0398), L2-LKP (p=0.0294) and L2-LKPR (p=0.0249) groups developed significantly higher atypical cells (Figure 6F). *Kras* altered organoids also showed significantly higher proliferation and DNA damage in the epithelial cells (Figure 6G-H). This data mimics the in-vivo findings and affirms that a *Kras* mutation is important, however it needs inactivation of at least one tumor suppressor to induce dysplastic *in-vitro* growth.

**Figure 6:**
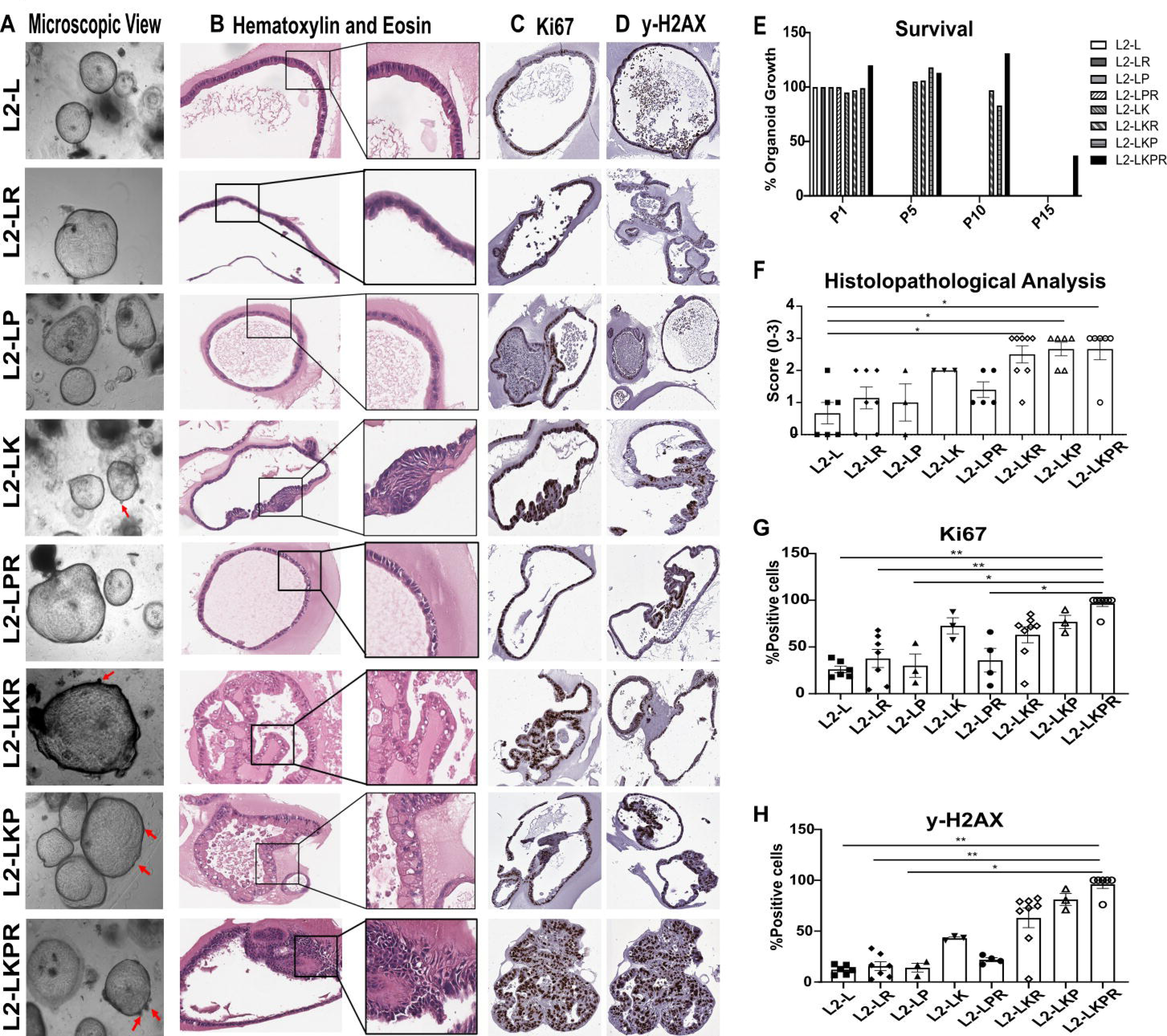
3D organoids mimic the epithelial characteristics of the in-vivo model. Representative images of organoids from different mice genotypes, (A) Microscopic images (Scale bar 200μm), (B) Histology (Scale bar 50 μm and 100μm), (C) Proliferation (Scale bar 50μm), (D) DNA damage (Scale bar 50μm), (E) Survival growth of organoids from passage 1-15 (n=3-8), (F) Statistical analysis of the histology of organoids (L2-L vs. L2-LKR, *p=0.0398; L2-L vs. L2-LKP, *p=0.0294; L2-L vs. L2-LKPR, *p=0.0249; n=3-8), (G) Immunohistochemistry for Ki67 (L2-L vs. L2-LKPR, **p=0.0015; n=3-8), and (H) Immunohistochemistry for y-H2AX (L2-L vs. L2-LKPR, **p=0.0013; n=3-8). Statistical analysis was done using Kruskal-Wallis test in addition to adjusted p-value with Dunn’s multiple comparison test.

### Genetic and epigenetic modifications occur with increased in mutational load in the mouse model

The distinct phenotypes of the different mouse models resulted in different survival rates (Suppl. Figure 4A). Mice with single mutations (L2-LP, L2-LR and L2-LK) survived longer compared to mice with double (L2-LPR > L2-LKR > L2-LKP) or triple mutations (L2-LKPR, maximum 6 months). L2-LKP survived only 9-months, indicating that *Kras-Tp53* alteration was more tumorigenic compared to *Kras-Rb1* alteration.

Reduced survival rates correlated with increased chromosomal instability as analyzed by low coverage whole genome sequencing and determination of genetic copies based on copy number variations (CNV). As shown previously, no chromosomal rearrangements were observed in L2-IL-1b mice [16] (Suppl. Figure 4B, C). However, CNV determination of dysplastic GEJ tissue from the various mutant mouse lines, showed increasing chromosomal aberrations with the accumulation of genetic alterations (Suppl. Figure 4B-F), both tumor and organoids from L2-LKPR mice (Suppl. Figure 4E, F) compared to L2-L (Suppl. Figure 4C, D).

Since the *KrasG12D* mutation was the driving carcinogenesis in the L2-LKPR mouse model, we assessed if *Kras* amplification also occurred in the murine GEJ tissue (Suppl. Figure 4G) and organoid system (Suppl. Figure 4H). Indeed, an amplification of the *Kras* gene could be detected in L2-LK, L2-LKR, L2-KP and L2-LKPR cohorts in-vivo and in-vitro. But no *Kras* amplification was observed in L2-LP, L2-LR and L2-LPR cohorts (Suppl. Figure 4G-H).

Loss of the tumor suppressor *p16^INK4A^* is mainly due to epigenetic silencing, e.g. promoter hypermethylation and reduced levels of *p16* are indicative of a poor outcome. To determine, if this also occured during cancer development, methylation of the *p16* gene[25] was determined for the various genotypes in-vivo and in-vitro. No changes in the *p16* methylation levels in GEJ tissue was observed (Suppl. Figure 4I), likely due to a heterogenous mix of cell types including epithelial, stromal, and immune cells within the sample. Testing the more homogenous epithelial cell population of the organoids revealed a gradual increase in *p16* methylation with increasing mutation load (Suppl. Figure 4J).

### Targeting *Kras* signaling abrogates esophageal tumor growth

Our mouse model clearly identified *KrasG12D* as the driver mutation of tumor growth. To targeted KRAS signalling and inhibit MAPK pathway activity, a combinatorial treatment was assessed (Figure 7A), combining a SHP2 inhibitor (SHP2i; RMC-4550) with a MEK inhibitor (MEKi; Trametinib) or with an ERK inhibitor (ERKi; LY3214996) as described previously[12, 26]. First, we analysed the inhibitors’ ability to inhibit MAPK pathway activity in murine and human organoids expressing mutant *Kras*. A range of different concentrations of SHP2i-MEKi and SHP2i-ERKi inhibitor combinations was tested (Suppl. Figure 5A and B), indicating that SHP2i (RMC-4550; 5μM), MEKi (Trametinib; 6.25nM) and ERKi (LY3214996; 5μM) could synergistically inhibit KRAS-driven tumor formation. Next, we treated murine organoids with either monotherapy or dual drug therapy. Organoids isolated from L2-LKPR mice showed reduced growth characteristics and cell death when treated with SHP2i+MEKi or SHP2i+ERKi (Figure 7B). Importantly, the same mono- and dual therapy did not affect L2-LPR organoids, which remained viable until day 3 (Figure 7C). This indicated that the dual therapy was KRAS specific (Figure 7D) and that the combination therapy was more effective compared to the monotherapy (Figure 7D). To assess MAPK activity in response to the different treatments the protein levels for SHP2, ERK, RSK1 and AKT and for phosphorylated pSHP2, pERK, pRSK1 and pAKT were determined. Reduced expressions of phosphorylated proteins was observed after the combination treatment (SHP2i+MEKi and SHP2i+ERKi) compared to the monotherapy (SHP2i) and the untreated groups or samples that did not express mutant *Kras* (Figure 7E, Suppl. Figure 6).

**Figure 7:**
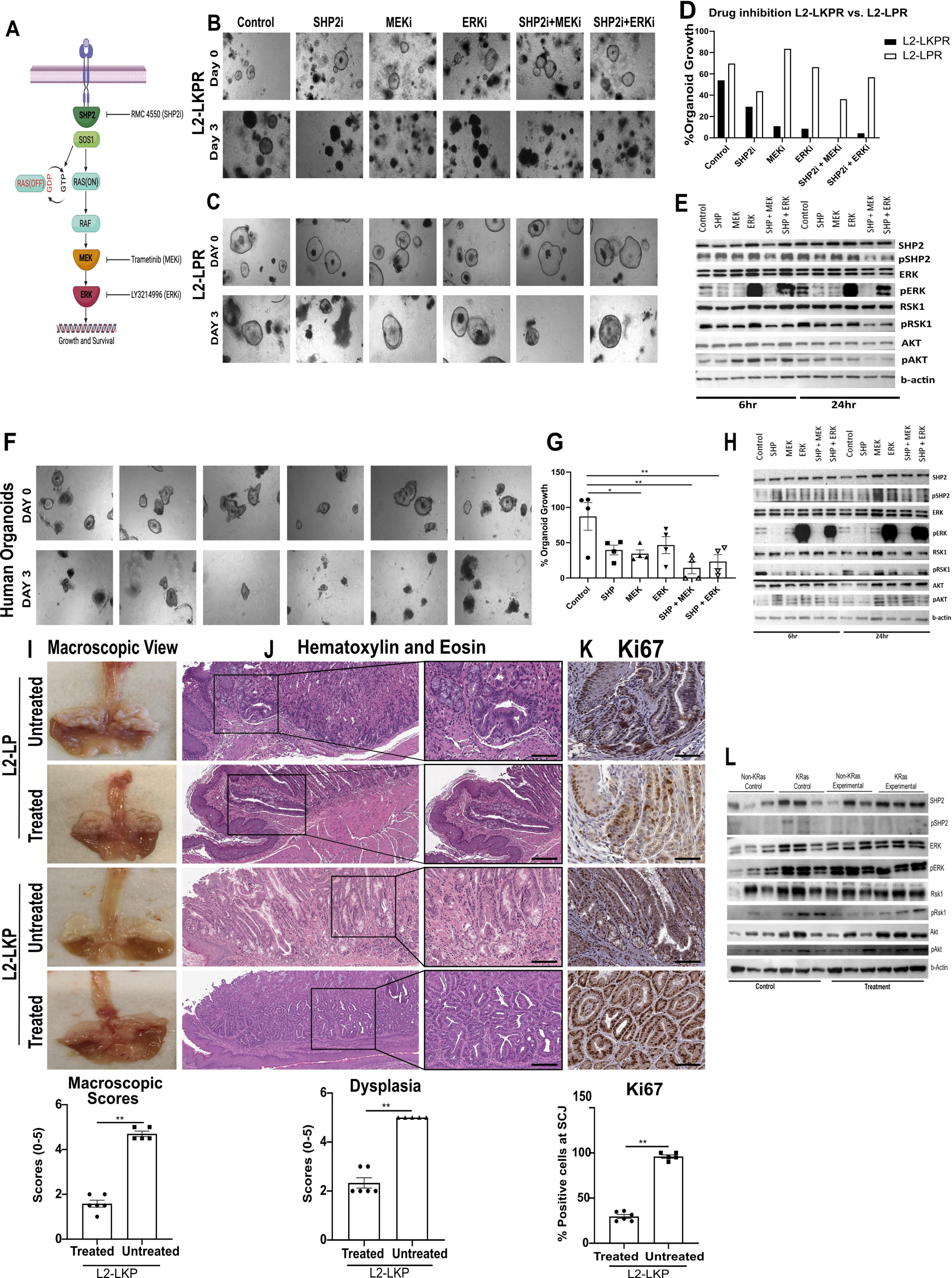
Dual-targeted therapy abrogates *Kras*-mediated tumors in EAC. Representative images of comparison of murine organoids with control, monotherapy, and dual therapy in on day 0 and 3 (B) L2-LKPR organoids and (C) L2-LPR organoids; (D) Comparison of organoid growth after drug treatment in L2-LKPR vs. L2-LPR organoids (*p=0.0313; n=3-5; paired two-tailed t-test); (E) Immunoblot with lysates collected at 6hr and 24hrs from *Kras* organoids (L2-LKPR) treated with monotherapy and dual therapy using the specific antibodies as indicated. b-actin was used as a loading control; (F) Representative images of human organoids with monotherapy and dual therapy on day 0 and 3. (G) Change in the growth of human organoids after drug treatment at day 3 (Control vs. MEK, *p=0.0409; Control vs. SHP+MEK, **p=0.0031; Control vs. SHP+ERK, **p=0.0099; n=4); (H) Immunoblot with lysates collected at 6hr and 24hr time-points from human organoids. Comparative analysis of (I) Macroscopic tumor (**p=0.0022; n=5-6); (J) Histopathology (**p=0.0022; n=5-6); (K) Proliferation (**p=0.0043; n=5-6) of in-vivo dual SHP2i+MEKi therapy in treated and untreated L2-LP and L2-LKP mice cohorts; (L) Immunoblots on lysates from mice tissue (n=3 per group). One experiment each was performed for immunoblots for mice and human organoids and mice tissue. Full scan images are shown in Suppl. Figure 5. For H-J, Mann-Whitney t-test was performed.

To translate these findings, we treated human organoids generated from GEAC endoscopic biopsies samples (BarrettNET registry)[27]. Prior to treatment a *KRAS* amplification was confirmed (Suppl. Figure 3C). Dual inhibitor combination with SHP2i+MEKi (**p=0.0031) and SHP2i+ERKi (**p=0.0099) resulted in significantly decreased organoid growth (Figure 7F, G) compared to the single inhibitors or control groups. Protein expression analysis showed a decrease in the phosphorylated proteins for dual combinations compared to monotherapy and control groups and confirmed a specific KRAS signalling inhibition (Figure 7H, Suppl. Figure 6).

Finally, we test the combination treatment SHP2i (RMC-4550) plus MEKi (Trametinib) in L2-LP and L2-LKP mice. As previously described,[26] mice received the combination of 1mg/kg Trametinib and 10mg/kg RMC-4550 drugs for 14 days via oral gavage (Suppl. Figure 5D). The dual drug treatment resulted in a significant decrease in macroscopic tumor growth at the GEJ in treated L2-LKP mice with 67% (4/6) of mice with ≤1mm of tumor size compared to untreated L2-LKP mice (Figure 7I) or treated L2-LP mice (Suppl. Figure 5E). Histopathological evaluation confirmed a significant treatment response in treated L2-LKP mice with 67% (4/6) showed reduced dysplasia with few atypical cells compared to untreated L2-LKP mice where 100% (5/5) mice developed invasive cancer (Figure 7J). This correlated with reduced proliferation (Figure 7K) at the GEJ of treated vs. untreated L2-LKP mice. Protein expression analysis confirmed the treatment effect on the MAPK pathway with reduced expressions of phosphorylated proteins in the treated compared to untreated cohorts (Figure 7L). In conclusion, our mouse and human organoid as well as preclinical data suggest that targeting KRas signalling emerges as an effective treatment option in EAC.

## Discussion

GEAC pathogenesis involves the proximal expansion of cardia progenitors, due to an inflammatory microenvironment[1, 5, 28]. A high mutation rate and clonal complexity of BE is evidence of an evolutionary process within progenitor cell populations that begins long before the development of a detectable malignancy or metaplasia[2]. Utilizing the L2-IL-1b mouse model[18] we demonstrate that inactivation of *Tp53* or *Rb* in such progenitor cells only mildly affects the dysplastic phenotype, but rather accelerates metaplasia development. In contrast, a *Kras* mutation had a profound effect on the development of dysplasia and when combined with mutated *Tp53* or *Rb* led to tumorigenesis. *LGR5* in this model is used to label a set of cells that includes a cell of origin for GEAC, affirming the understanding that EAC and GC originate from similar gastric progenitor cell population[1, 5].

Although *Tp53* is preferentially altered in patients with nondysplastic BE, which progress to cancer, in our mouse model (L2-LP) with conditional knockout of *Tp53*, we observed only a mild acceleration of the phenotype, suggesting that additional factors or different alterations are needed to induce cancer. Complete loss of *Tp53* in *Lgr5* cells did not induce other chromosomal alteration, as demonstrated in CNV analysis in organoids and GEJ tissue. Nevertheless, *TP53* is most commonly mutated in dysplastic BE patients, and we did not study a *Tp53* mutated model, which could have a different outcome[29]. *Rb1* is downstream of *p16^INK4A^* in the CDKN2A signalling pathway, and while we disrupted p16 signalling by conditional knockout of *Rb1*, this on its own was insufficient to induce invasive cancer or chromosomal aberrations or alter organoid growth. *CDKN2A* can be mutated[6] or epigenetically altered in BE[9] and we observed more profound metaplasia development upon loss of *Rb* and *Tp53*, suggesting that these alterations in *LgR5+* cells lead to a more differentiated phenotype with reduced progression to dysplasia. Interesting it was shown that *LgR5+* cell activation was important in directing gland reconstitution during inflammation,[30, 31] and previous analysis of GEAC cases could correlate *RB1* methylation with intestinal metaplasia.[32] We would argue that the observed p21 activation induced cell cycle arrest in *Tp53* or *Rb1* altered cells. Cell cycle arrest might correlate with increased differentiation to metaplasia and thus p21 expression in combination with differentiation markers, such as goblet cell differentiation or Notch signalling.[16, 33, 34] Recently, a physiologic role of somatic mutations in preserving tissue homeostasis during repeated damages was described in pancreatic carcinogenesis, where *KRAS* mutations appeared beneficial during chronic inflammation[35]. Oncogene mediated metaplasia might thus be an adaptation to tissue damage under chronic inflammation as seen in GERD and BE patients and as seen here for *Rb1* or *Tp53* in our mouse model. We reported early on the regenerative capability of progenitor cells during chronic inflammation,[2, 36] and the ability to sustain an adaptive response would represent an evolutionary advantage, with the development of metaplasia (BE) representing a protective epithelial condition in the setting of esophagitis, but with the inherent risk of progression to neoplasia following additional oncogenic alterations in defined progenitor cells.

Upon combined deletion of *Tp53* and *Rb1*, the mice developed high-grade dysplasia. Indeed, in gastric cancer co-deletion of *CDKN2A* (upstream of *RB1*) and *TP53* sensitizes to DNA damage blockade and induces premalignant lesions[37, 38, 39]. Given the expansion of clones with altered *TP53* appear within *CDKN2A* deficient clones in BE,[39] we would hypothesize that loss or alteration of *Tp53* confers a selective advantage in cells of origin and therefore is an early driver of progression to GEAC, if a second hit is achieved. It was similarly shown that deletion of *Tp53* in gastric cells conferred a selective advantage and promoted gastric dysplasia in combination with carcinogens.[40]

A third – so far not widely recognized - genetic alteration that appeared crucial in 15% of GEAC is a *KRAS* mutation or amplification. Indeed, *Kras* mutation induced high-grade dysplasia, suggesting a more powerful oncogenic potential than *Tp53* or *Rb1*. Upon introduction of *Kras* mutation together with either *Tp53* (L2-LKP) or *Rb1* (L2-LKR) we observed invasive carcinoma, which was further accelerated with all three alterations in L2-LKPR mice, resembling human tumors. Of note, the development of invasive cancer correlated with downregulation of Tp53 and p21 signalling. As increased chromosomal instability and epigenetic modifications also supported the intrinsic nature of the mutations, it is important to note that chromosomal instability could only be achieved with the combination of genetic alterations, suggesting that at least two if not three events are necessary in GEAC to induce tumorigenesis.

The specific mechanisms of the tyrosine phosphatase SHP2 (SH2 domain-containing phosphatase 2) in RAS and MAPK signalling have been an area of active study in the RAS field.[26, 41, 42] It has previously been demonstrated that loss of SHP2 function leaves *Kras*-mutant cancer cells uniquely sensitive to MEK and ERK inhibition.[26, 43] *Kras*-amplified GEAC models were insensitive to MAPK blockade and inhibition of SHP2 attenuated this adaptive process to MEK inhibition.[12] Of note, earlier characterization reported the occurrence of frequent KRAS amplification in GEAC and concluded that the increased protein expression, likely aided by the lack of canonical somatic mutations that alter the KRAS equilibrium, creates a dynamic state with greater potential to mobilize KRAS-GTP.[12, 13] Indeed, *Kras* mutation in the mouse model correlated with increased amplification, supporting *Kras* mutations as general model for *Kras* activation (such as with amplification). We would hypothesise that the frequently seen *KRAS* amplification in GEAC is secondary to inflammation induced KRAS pathway activation that might lead to fixed *KRAS* amplification. Such amplification in addition to other early alterations such as Tp53 would overcome growth arrest and finally promote cancer development. Similarly, an evolutionary dose dependent activation of mutant *KRAS* has been described in pancreatic carcinogenesis.[44]

MEK and ERK inhibitors have failed or not yet been approved for clinical use in GEAC, which is partly due to rapidly occurring resistance.[45] We show that only combinatorial pharmacologic targeting of ERK or MEK signaling with a SHP2 inhibitor prevents human and mouse organoid growths in a KRAS dependent manner. The combination of these inhibitors upstream and downstream of RAS significantly exceeded the effects of each alone and was strongest with the SHP2/MEK inhibitor combination. In our mouse model tumor development and growth was abrogated in *Kras* altered mice, suggesting that SHP2 inhibition was *Kras* dependent. Thus, our data indicate a clinical utility of dual SHP2/MEK inhibition as a targeted therapy approach for *Kras*-altered GEAC. Further clinical studies are needed, especially as previously inhibition of SHP2 could attenuate tumor progression in both in vitro and in vivo settings.[12]

In conclusion, with the generation of the L2-LKPR mouse model, we recapitulated human carcinogenesis with a first murine invasive GEAC model and show that *KRAS* is an important driver of tumorigenesis, at least in combination with a second genetic alterations. Targeting KRAS alterations could be important for GEAC and warrants confirmation in clinical studies, especially as specific KRAS inhibitor are on the horizon.[46, 47] In addition, a concept of single genetic alteration inducing oncogene mediated metaplasia as an adaptation to tissue damage under chronic inflammation as seen in GERD and BE might emerge as an important factor for patients under surveillance for GEAC.

