## Supplemental Information for "KRas, in addition to Tp53 is a driver for early carcinogenesis and a molecular target in a mouse model of invasive gastro-esophageal adenocarcinoma"

### Supplementary Experimental Procedures:

#### Tissue preparation and disease evaluation

For mouse samples, after euthanasia, stomach and esophagus tissues were fixed in 4% paraformaldehyde and paraffin-embedded. Slides were stained with H&E (Haematoxylins and eosin) and evaluated by a mouse pathologist as described previously [1].

For the disease progression assessment, macroscopic scoring was performed. Each stomach and esophagus was evaluated for tumor coverage, individual tumor size, total tumor size, and summed for an overall macroscopic score as previously described [1, 2]. Histopathological scoring was performed by an experienced mouse pathologist using the previously established criteria [3]. Inflammation was scored by the percentage of different immune cells (mostly neutrophilic and myeloid cells) in a defined tissue area of the GEJ in a high-power field evaluation. Metaplasia was evaluated by the abundance of mucus producing or cells per gland and the abundance of glands with mucus producing cells in the BE area at the GEJ. Dysplasia was evaluated by the amount of cellular atypia and the presence of low- or high-grade dysplasia in single or multiple glands. Invasive carcinoma was evaluated on the clusters of undifferentiated cells present in epithelial layers.

#### Immunohistochemistry and TUNEL Assay

Standard immunohistochemical procedures with citrate buffer heat-mediated antigen retrieval (H-3300, Vector labs) were performed using the following antibodies: rabbit anti-Ki67 antibody (Abcam #ab15580, 1:1000, 4°C overnight), rabbit anti-aSMA (Abcam #ab5694, 1:400, 2h room temperature), rabbit anti-γ-$H_2AX$ (Cell Signalling #9718 1:750, 4°C overnight). EDTA buffer antigen retrieval was performed for the following antibodies: rabbit anti-p53 (Leica #NCL-L-p53-CM5p, 1:150, 4°C overnight) and p21 (Abcam #ab107099, 1:350, 4°C overnight) for mouse tissue. Quantification was accessed as percentage of positive cells or areas in BE regions. In situ Cell Death Detection Fluorescein kit (Roche, Germany) was used to perform TUNEL assay as described in the kit protocol.

**Three-dimensional organoid culture**

Organoids were processed from both the GEJ of mouse stomach [4] and the human patient BE/EAC biopsies resected during endoscopic procedures. Organoids were passaged every 7 days until growing healthy. Organoids were fixed with 4% PFA and were embedded in paraffin for histology as previously described [5]. The histology of organoids was graded on an ascending scale, from 0-3, based on the structure of the cells and the placement of the nucleus: 0, squamous epithelium; 1, columnar epithelium; 2, apoptotic cells and 3, undifferentiated cells. Ethical confirmation for use of Oragnoids was provided bei TUM Ethikkommisson No: 21-1380

**Methylation analysis**

For both methylation and genomic analysis, DNA was extracted from both tissue and organoids using the AllPrep DNA/RNA kit (80224, Qiagen) by following the instructions in the kit protocol.

For pyrosequencing 200ng of gDNA was processed using EZ DNA Methylation-Direct kit (Zymo Research, Irvine, USA) according to manufacturer's instructions. PCR amplification was performed using PyroMark PCR kit (Qiagen) using the primers: p16Ink4a-F (AGTAGTGTTTTTAGGGGTGT); p16Ink4a-R ((Btn)-CCATACTACTCCAAATAACTCTC) and p16Ink4a-Seq (GGAAGGAGGGATTTATTG). PCR products were pyrosequenced using PyroMark Q48 Advanced CpG reagents on a PyroMark Q48 Autoprep instrument (Qiagen). For assay optimisation, methylated (100%), non-methylated (0%) and a scale of control samples with the following DNA methylation: 25%, 50% and 75% were used. The methylated control was prepared using the CpG methyltransferase enzyme (M.SssI; Thermo Scientific). The non-methylated control was prepared using the REPLI-g Mini kit (Qiagen).

**Genomic Analysis**

LcWGS and Amplicon sequencing was performed on a NextSeq 500 (Illumina) in a 75 cycles single end run. Amplicon sequencing was used to assess *Kras* amplification. The raw reads were mapped to *Kras* reference sequence (Ensemble release GRCm38p4, Genome Reference Consortium) and variant allele frequencies on chromosome 6 at position 145246771 were calculated.

**Western Blotting**

Tissues were immediately snap frozen in liquid nitrogen at the time of harvest. Tissue or cells were lysed in RIPA lysis buffer supplemented with protease and phosphatase inhibitors and protein concentrations were determined by Bradford assay (Bio-Rad). Proteins were separated by SDS-PAGE in Laemmli buffer, transferred to 0.2µm nitrocellulose membrane and detected with the following antibodies: ERK1/2 (4307), pERK1/2 (9101), SHP2 (3397), AKT (9272), pAKT (9271) and pRSK1 (9344) were purchased from Cell Signalling. pSHP2 (ab62322) was from Abcam. RSK1 (PA5-29215) was from Invitrogen and β-actin-HRP (A3854) purchased from Sigma Aldrich served as the loading control. Signal detection was performed using HRP-conjugated secondary antibody and an enhanced chemiluminescent reagent (Amersham, GE Healthcare). The signals were visualized by BioRad Gel Documentation system.

**Drugs and Inhibitors**

MEKi (Trametinib), ERKi (LY3214996) and SHP2i (RMC-4550) were purchased from Selleckchem. The drugs were dissolved in DMSO to yield 5-50mM solutions and were stored at -80°C.

**In-vitro drug screening and treatment**

Organoids were cultured and expanded from murine GEJ tissue and patient BE/EAC biopsies using previously described protocols [4]. For high-throughput drug screening, single cell suspensions were obtained from organoids using mechanical and enzymatic degradation. Cell-Matrigel suspensions were plated in 96-well plates ($1X10^3$ cells per well). After 24 hours, titration treatments were initiated, and the cell viability was measured after 72 hours via CellTitre-Glo 3D Viability Assay (Promega) luminescence on a FLUOstar OPTIMA microplate reader (BMG Labtech).

Both murine and human organoids were treated with 5 µM SHP2i, 5 µM ERKi, 6.25nM MEKi and a combination of SHP2i-ERKi and SHP2i-MEKi. Organoids were collected for protein isolation at 6hr and 24hr time-points.

**Quantitative analysis of drug synergy**

Drug synergy was calculated using CompuSyn software (version 1.0) This software generates combination index (CI) values, where CI<0.75 indicates synergism; CI=0.75-1.25 shows additive effects and CI>1.25 indicates antagonism.

**In-vivo therapy treatment**

For in-vivo treatments, a combination of Trametinib and RMC-4550 was administered via oral gavage in L2-LP and L2-LKP mice cohorts. All the mice received drugs for 5 days, 2 days off for two weeks. Drugs were prepared as described previously [6].

**Statistical Analysis**

Each data point on every graph represents one mouse. For comparison between two groups, Mann-Whitney t-test was used. For comparison between more than two groups, one-way analysis of variance (ANOVA) with Dunn's multiple comparison test was used. Data is represented as mean ± standard error of mean (SEM) (****p<0.0001, ***p<0.001, **p<0.01, *p<0.05). All the statistical analysis was performed with GraphPad Prism 8.0.2 software.

### Supplementary Table

**Supplementary Table 1:** List of all the different genotypic combinations with their respective annotations

### Supplementary Figures

**Supplementary Figure 1: Mutational frequencies based on public data** Illustration of mutational frequencies of *TP53, CDKN2A, KRAS* and *RB1* from the publicly available data for the respective genetic alterations [7, 8].

**Supplementary Figure 2: Rb knock-out has an accelerating effect on metaplasia.** (A)Representative images of single transgenic groups L2-L, L2-LR, L2-LP and L2-LK indicating in black arrows the presence of metaplastic cells at the SCJ, (B) Quantification of metaplasia formation in (L2-L vs. L2-LKR+/-, ****p<0.0001; L2-L vs. L2-LR, *p=0.0229; L2-L vs. L2-LP+/-, p=0.3130; L2-L vs. L2-LP, p=0.3972; L2-L vs. L2-LK, *p=0.0359; L2-L vs. L2-LPR, **p=0.0040, n=6-14). For statistical analysis, Kruskal-Wallis test was performed in addition to adjusted p-value with Dunn's multiple comparison test. Groups with grey backgrounds are the same mice cohorts as their respective groups in Figure 1, 2 and 3.

**Supplementary Figure 3: *Kras* in combination with heterozygous *Tp53* and *Rb1* has the potential to develop invasive cancer.** (A) Macroscopic overview of tumor formation in L2-L (*L2-IL-1b.Lgr5+/-),* L2-LKR+/- (*L2-IL-1b.Lgr5+/-.Kras+/-.Rb1+/-),* L2-LKP+/- (*L2-IL-1b.Lgr5+/-.Kras+/-Tp53+/-),* L2-LKP+/-R+/- (*L2-IL-1b.Lgr5+/-.Kras+/-.Tp53+/-.Rb1+/-),* L2-LKP+/-R-/- (*L2-IL-1b.Lgr5+/-.Kras+/-.Tp53+/-.Rb1-/-),* L2-LKP-/-R+/- (*L2-IL-1b.Lgr5+/-.Kras+/-.Tp53-/-.Rb1+/-)* mice; (B) Quantification of macroscopic scores (L2-L vs. L2-LKR+/-, ***p=0.0008; L2-L vs. L2-LKP+/-, *p=0.0372; L2-L vs. L2-LKP+/-R+/-, p=0.0648; L2-L vs. L2-LKP+/-R-/-, ****p<0.0001; L2-L vs. L2-LKP-/-R+/-, **p=0.0011; n=7-16); (C) Distribution of mice by macroscopic scores from B; (D) Representative images of hematoxylin and eosin staining of L2-L, L2-LKR+/-, L2-LKP+/-, L2-LKP+/-R+/-, L2-LKP+/-R-/-, L2-LKP-/-R+/- mice to assess the histopathology, scale bars 100µm and 200µm; (E) Inflammation (L2-L vs. L2-LKR+/-, *p=0.0169; L2-L vs. L2-LKP+/-, ***p=0.0004; L2-L vs. L2-LKP+/-R+/-, *p=0.0312; L2-L vs. L2-LKP+/-R-/-, **p=0.0045; L2-L vs. L2-LKP-/-R+/-, **p=0.0029; n=7-14); (F) Metaplasia (L2-L vs. L2-LKR+/-, ***p=0.0006; L2-L vs. L2-LKP+/-, **p=0.0013; L2-L vs. L2-LKP+/-R+/-, ****p<0.0001; L2-L vs. L2-LKP+/-R-/-, ****p<0.0001; L2-L vs. L2-LKP-/-R+/-, ***p=0.0003; n=7-14); (G) Dysplasia (L2-L vs. L2-LKR+/-, **p=0.0032; L2-L vs. L2-LKP+/-, p=0.7801; L2-L vs. L2-LKP+/-R+/-, **p=0.0054; L2-L vs. L2-LKP+/-R-/-, ***p=0.0008; L2-L vs. L2-LKP-/-R+/-, ***p=0.0006; n=7-14); (H) Distribution of mice by dysplasia scores from G; (I) Representative images of immunohistochemistry for Ki67 for L2-L, L2-LKR+/-, L2-LKP+/-, L2-LKP+/-R+/-, L2-LKP+/-R-/-, L2-LKP-/-R+/- mice, scale bar 50µm; (J) Immunohistochemistry for Ki67 (L2-L vs. L2-LKR+/-, **p=0.0085; L2-L vs. L2-LKP+/-, p>0.999; L2-L vs. L2-LKP+/-R+/-, **p=0.0014; L2-L vs. L2-LKP+/-R-/-, ****p<0.0001; L2-L vs. L2-LKP-/-R+/-, ***p=0.0001; n=6-14); For statistical analysis, Kruskal-Wallis test was performed in addition to adjusted p-value with Dunn's multiple comparison test. Groups with grey backgrounds are the same mice cohorts as their respective groups in Figure 1.

**Suppl. Figure 4: Genomic and epigenomic instability accelerates with increase in mutational burden.** Survival curves of the homozygous groups compared to the control L2-L group in months (A, left side) Single mutations (n=7-13) and (A, right side) Multiple mutation combinations (****p<0.0001, n=6-16). Statistical representation of the chromosomal aberrations in epithelial cells between different mutational alterations with dysplasia score ≥ 4 (B) Number of genes based on Copy Number Variations (L2-L vs. L2-LKR, p=0.6209; L2-L vs. L2-LKP, p=0.3291; L2-L vs. L2-LKPR, p=0.1430; n=3-15). Representative images of the chromosomal instability in L2-L group in mice (C) Tissue and (D) Organoids compared to the L2-LKPR group in mice (E) Tissue and (F) Organoids. Comparison of KRasG12D amplification of control, single, double, and triple transgenic mice cohorts in (G) Tissue (*p=0.0042; n=3) and (H) Organoids (**p=0.0042; n=3). Methylation analysis of control, single, double, and triple transgenic mice cohorts in (I) Tissue (p=0.5326; n=6) and (J) Organoids (p=0.1573; n=3). Statistical analysis was done using Logrank test for graph A and for all the other analysis Kruskal-Wallis test in addition to adjusted p-value with Dunn's multiple comparison test was performed.

**Supplementary Figure 5: Evaluation of drug dosage combinations in organoids and *Kras*-altered mouse model.** Drug combination synergy defined by combinatorial index in (A) Mouse organoids and (B) Human organoids; (C) Representative image of the LcWGS analysis of human organoid chromosome with a magnified image of chromosome 12; (D) Schedule for in-vivo dual drug treatment; (E) Histological analysis comparing control group L2-LP, treated vs. untreated on the parameters of macroscopic scores, dysplasia and immunohistochemistry of Ki67

**Supplementary Figure 6: Immunoblots** Full scan immunoblots from mouse and human organoid lysates and lysates from mice tissue as indicated in the figure.
