## Supplementary figures and images for "KRas, in addition to Tp53 is a driver for early carcinogenesis and a molecular target in a mouse model of invasive gastro-esophageal adenocarcinoma"

### Supplemental Figure 1

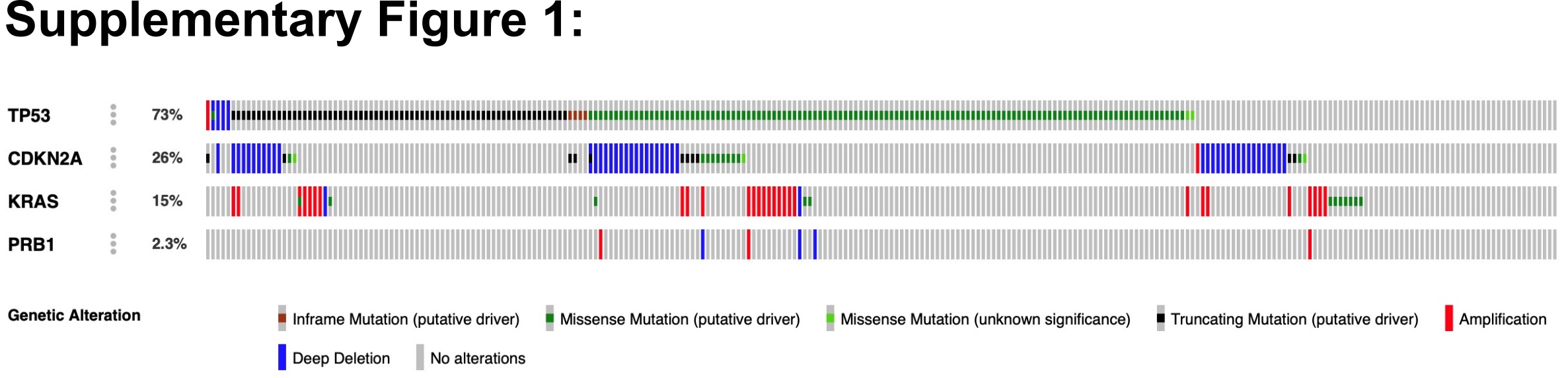

### Supplemental Figure 2

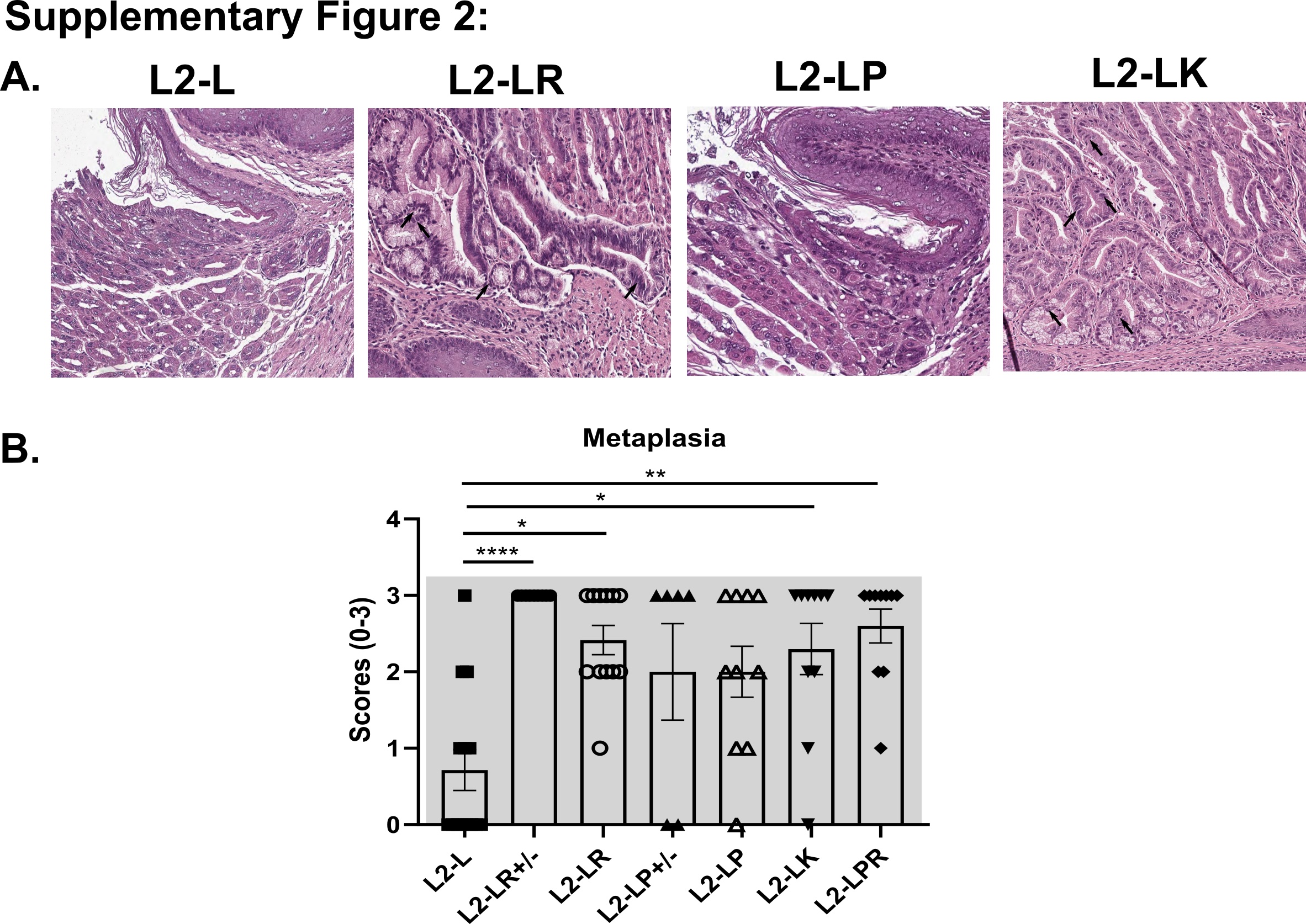

### Supplemental Figure 3

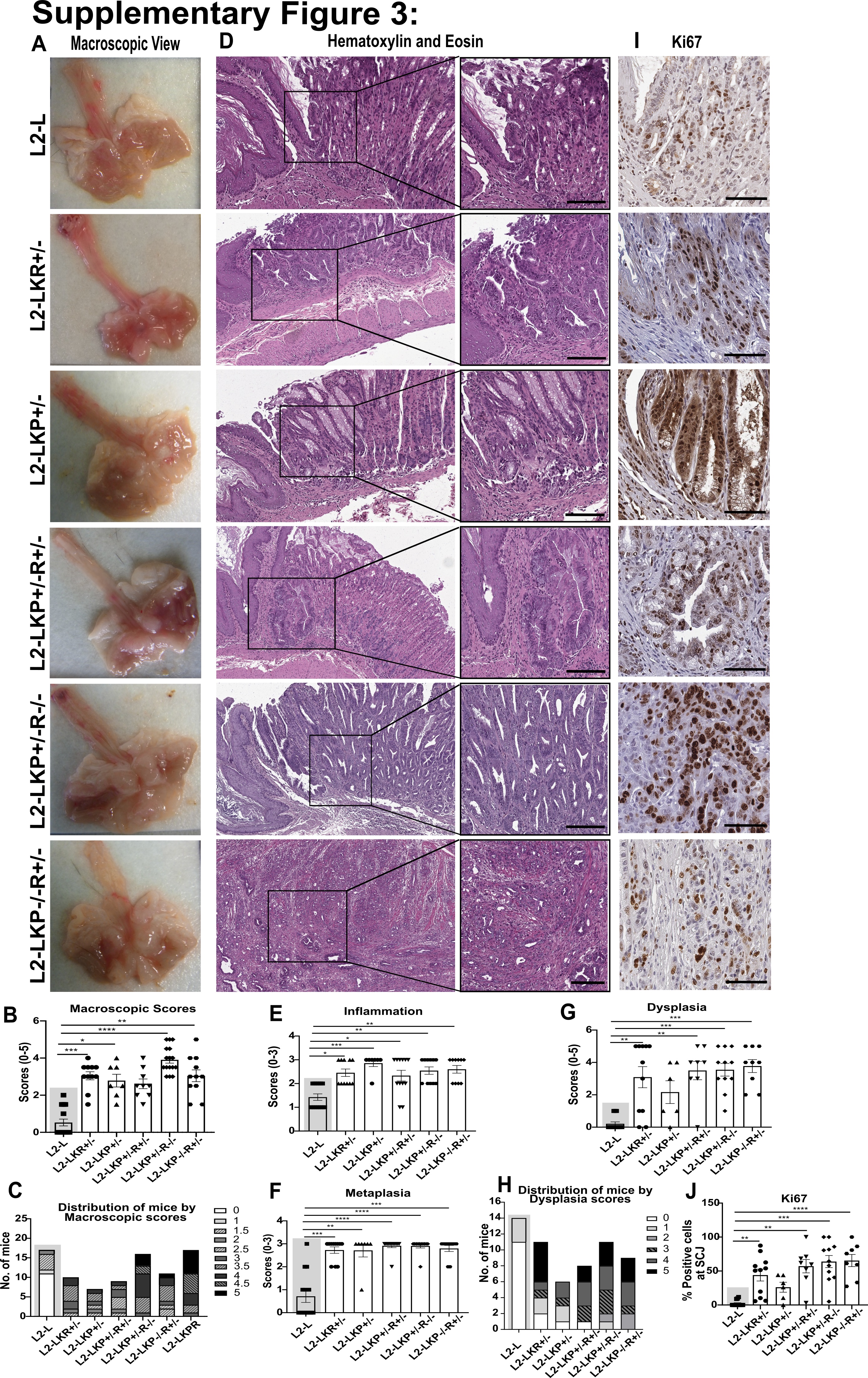

### Supplemental Figure 4

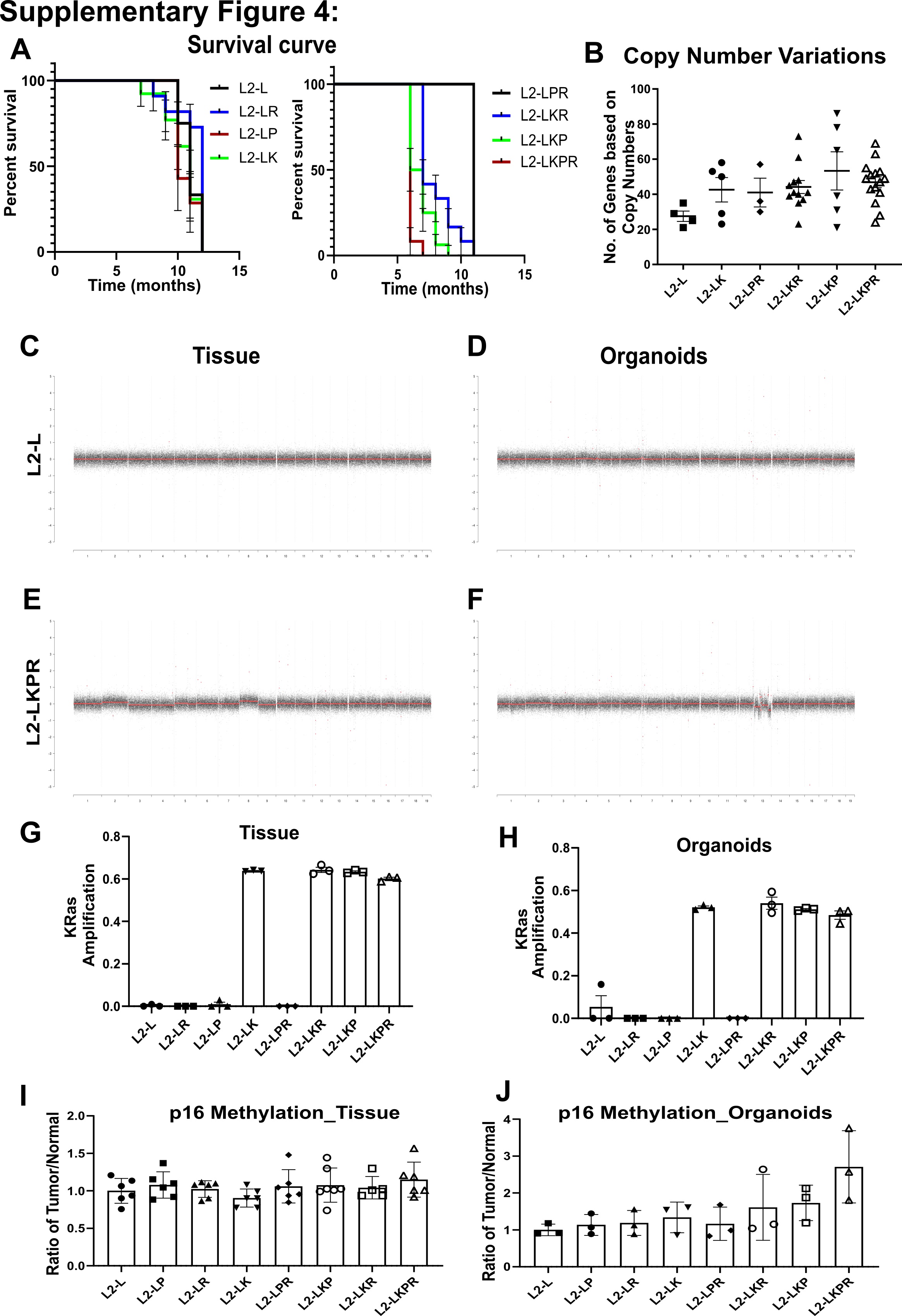

### Supplemental Figure 5

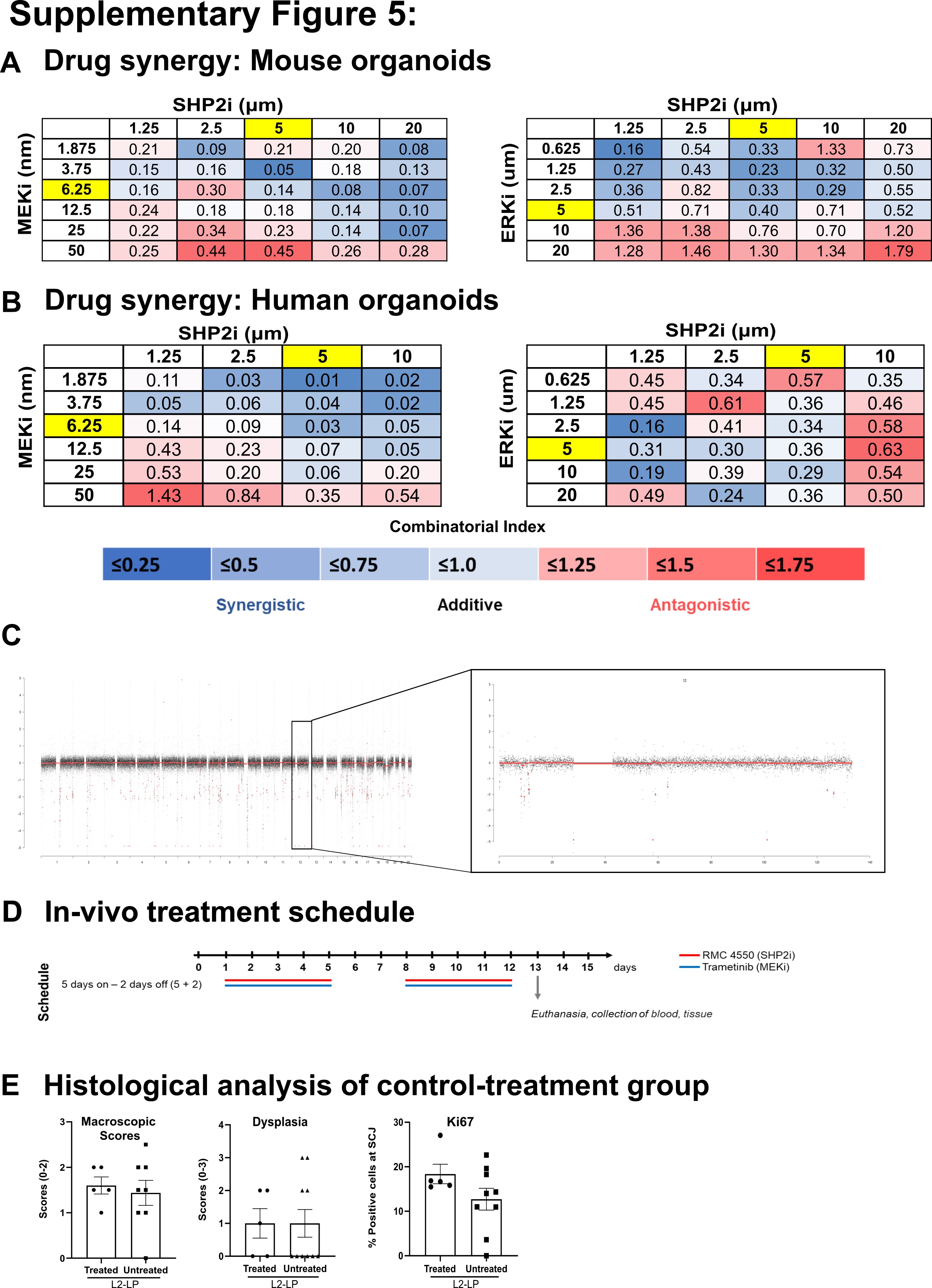

### Supplemental Figure 6

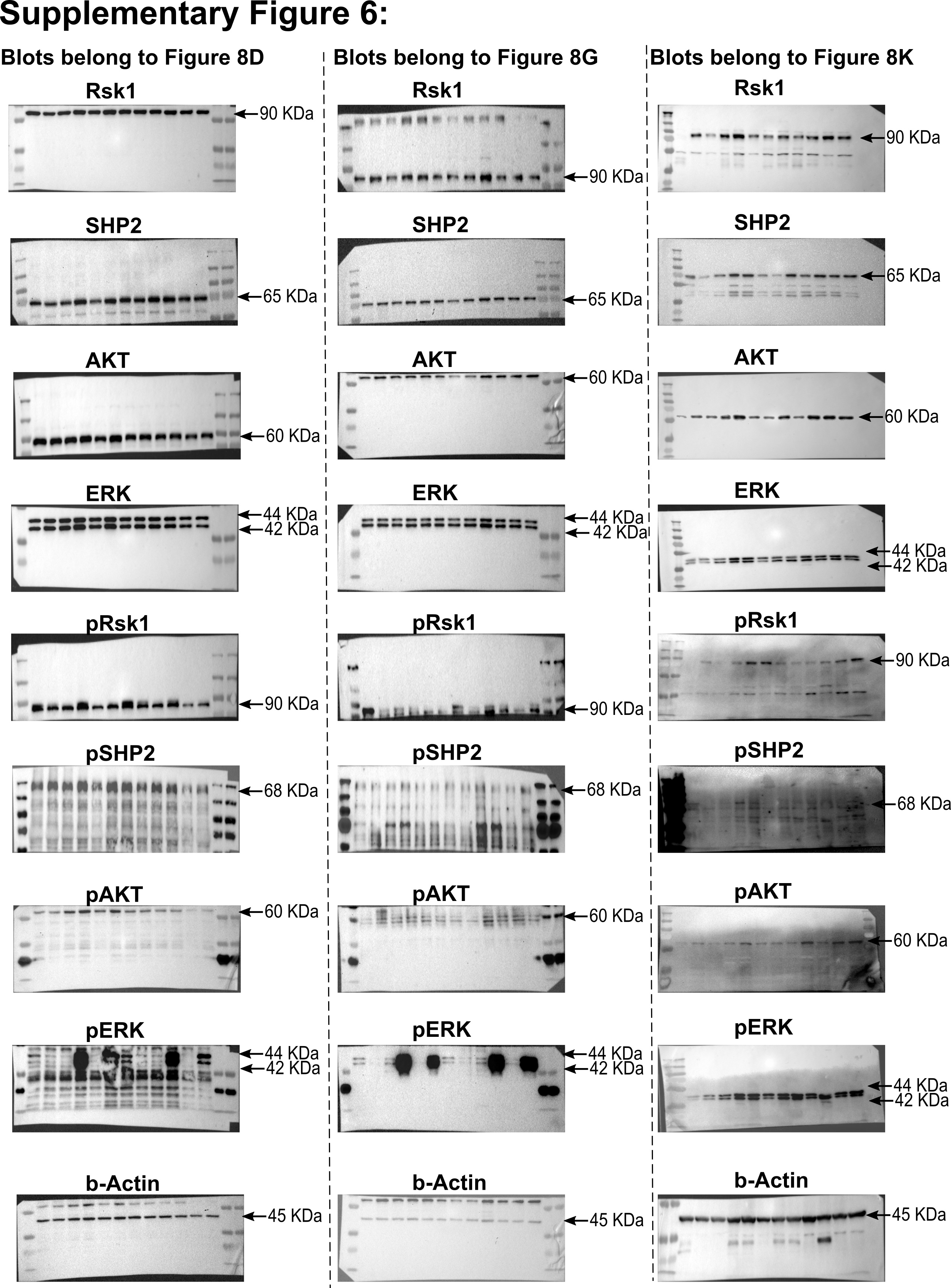

### Supplemental Table 1

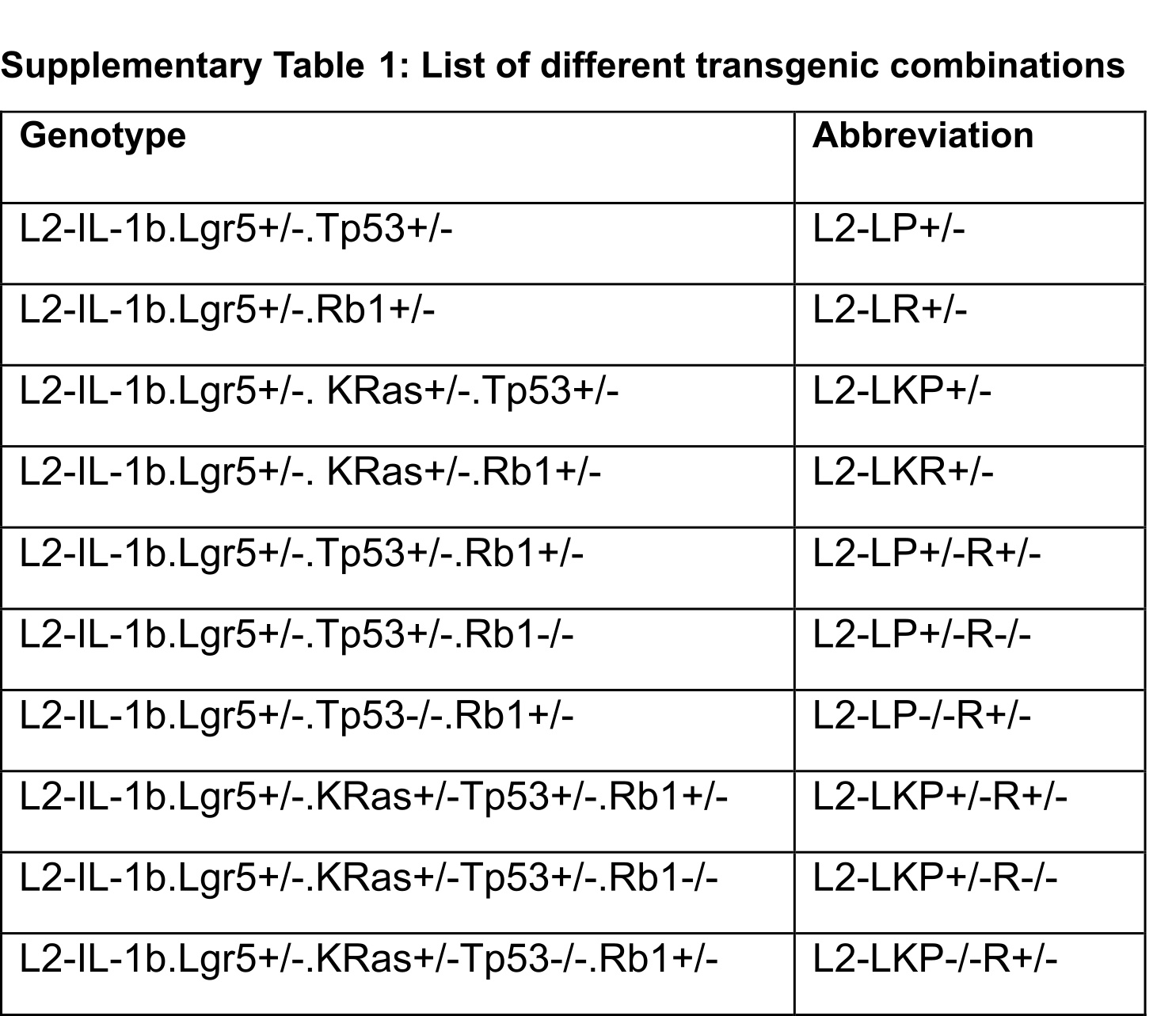
